# ATG13 dynamics in non-selective autophagy and mitophagy: insights from live imaging studies and mathematical modelling

**DOI:** 10.1101/370114

**Authors:** Piero Dalle Pezze, Eleftherios Karanasios, Varvara Kandia, Maria Manifava, Simon A. Walker, Nicolas Gambardella Le Novère, Nicholas T. Ktistakis

## Abstract

During autophagy, the ULK complex nucleates autophagic precursors which give rise to autophagosomes. We analysed by live imaging and mathematical modelling translocation of ATG13 (part of ULK complex) to autophagic puncta in starvation-induced autophagy and ivermectin-induced mitophagy. In non-selective autophagy, the intensity and duration of ATG13 translocation approximated a normal distribution whereas wortmannin reduced this and shifted to a log-normal distribution. During mitophagy, multiple translocations of ATG13, with increasing time between peaks were observed. We hypothesised that these multiple translocations arise because engulfment of mitochondrial fragments requires successive nucleations of multiple phagophores on the same target, and a mathematical model based on this idea reproduced the oscillatory behaviour. Significantly, model and experimental data were also in agreement that the number of ATG13 translocations is directly proportional to the diameter of the targeted mitochondrial fragments. Our data provide novel insights into the early dynamics of selective and non-selective autophagy.

## Introduction

Autophagy can be non-selective when the cargo is generic cytoplasmic material, providing the cell with a mechanism of nutrient supply for periods of starvation (Mizushima and Komatsu, 2011; Dunlop and Tee, 2014; Mony et al, 2016), or selective when it leads to the degradation of specific intracellular structures - damaged mitochondria, endoplasmic reticulum fragments, bacterial pathogens etc - forming the basis for a crucial cellular quality control system (Okamoto, 2014; Stolz et al, 2014; Randow and Youle, 2014; Anding and Baehrecke, 2017).

The starvation-induced pathway utilises a series of membrane re-arrangements after inactivation of the protein kinase complex mTORC1 (Saxton and Sabatini, 2017). This leads to the activation of the autophagy-specific protein kinase ULK complex formed by the protein kinase ULK1 (or its homologue ULK2), and the adaptors FIP200, ATG13, and ATG101 (Lin and Hurley, 2016; Zachari and Ganley, 2017). Once active, the ULK complex translocates to tubulovesicular regions of the ER which are characterised by the presence of vesicles containing the autophagy-specific ATG9 protein. On these sites the lipid kinase VPS34 complex I also translocates to produce PI3P and generate omegasomes from where autophagosomes spawn (Walker et al, 2008; Karanasios et al, 2016). In addition to this linear sequence, we have also proposed a PI3P-dependent positive feedback loop re-enforcing ULK binding to membranes in the early phases of omegasome-ULK expansion (Karanasios et al 2013). PI3P binds WIPI proteins, which in turn bind to the ATG16 protein, linking the formation of autophagosomal membrane to the complexes covalently modifying LC3/GABARAP - the protein binding autophagosome cargo - with phosphatidylethanolamine (Wilson et al, 2014; Yu et al, 2017).

In the case of selective autophagy, additional specific proteins (adaptors or receptors) are necessary to label the cargo and connect it with the forming autophagosome (Johansen and Lamark, 2011; Randow and Yule, 2014; Rogov et al 2014; Khaminets et al, 2015; Roberts et al, 2016). Triggering these adaptors or receptors leads to the recruitment of the LC3/GABARAP proteins, enabling the engulfment of the targeted membrane (Khaminets et al, 2016; Yamano et al, 2016). It should be noted however that, even in the absence of all LC3/GABARAP proteins, autophagic membranes in the process of engulfing cargo can be observed (Nguyen et al, 2016).

Mitophagy in tissue culture cells can be induced by a variety of methods including a number of pharmacological compounds, usually affecting the mitochondrial membrane potential or metabolite balance of the mitochondrial lumen (Georgakopoulos et al, 2017). In contrast to most of these drugs which require several hours to take effect, the anthelmintic lactone ivermectin (IVM) was also found to induce mitochondrial fragmentation and mitophagy within 30 minutes of treatment, making it an ideal tool in our recent study of mitophagy dynamics (Zachari et al, 2019). In that work, we observed that the translocation of the ATG13 protein to forming mitophagosomes occurred in several waves resembling oscillations (Zachari et al, 2019), something that we had not seen for the translocation of the same protein to starvation-induced autophagosomes. Here, we used complementary *in vitro* and *in silico* approaches to investigate the dynamics of ATG13 in the initiation of non-selective autophagy and ivermectin-dependent mitophagy. To be able to compare these two processes directly, we used spinning disk confocal microscopy to analyse the dynamics of the response, and the same quantitation protocols to follow the formation of ATG13-containing autophagy structures. Our imaging data were used to derive mathematical models of the two processes.

Data-driven mathematical modelling has been a valuable method for describing biological signalling networks and investigating novel regulatory mechanisms (Kholodenko et al, 2010; Le Novère, 2015; Tomlin and Axelrod, 2007) including autophagy dynamics (Martin et al, 2012).. Mathematical models have also been proposed for the upstream signalling of autophagosome formation (Szymańska et al, 2015; Dalle Pezze et al, 2016).

## Results

### A mathematical model for ATG13 dynamics during non-selective autophagy

To generate a mathematical model of the earliest visible accumulation of autophagic structures in non-selective autophagy, cells stably expressing GFP-ATG13 were imaged during starvation and data were collected in presence and absence of the PI3K inhibitor wortmannin (see also Karanasios et al, 2013). Representative time course images for one autophagy event are shown in **Figure 1A**. Wortmannin triggered a reduction of the ATG13 punctum intensity and lifetime, confirming the proposed PI3P-dependent positive regulation of ATG13 accumulation (Karanasios et al, 2013). Autophagy events occured in the cytoplasm asynchronously and the weak signal at the beginning of the aggregation makes it difficult to determine an exact starting time point. Therefore, we synchronised these data using the ATG13 maximum intensity peak (see **Materials and Methods** for details). We built a minimalistic model for the early steps of non-selective autophagy where ATG13 aggregates and dissociates following mass action reactions (**Figure 1B**). ATG13 accumulation and disappearance were non-linear, so we included a dependency for the process on the concentration of aggregated ATG13, with a partial reaction order *m*. To investigate the effect of wortmannin on these reactions, three models were investigated: 1) wortmannin regulating ATG13 aggregation, 2) wortmannin regulating ATG13 disappearance, and 3) wortmannin regulating both aggregation and disappearance. As ATG13 accumulation time courses were characterised by a rapid increase followed by a steep decrease, we tested two additional hypotheses. In the first, the switch between ATG13 accumulation and removal was regulated by an event happening after *t* seconds from the initiation of the process. This delay was estimated from the imaging data. In the second, the reactions of ATG13 accumulation and removal happened simultaneously without any additional regulation. The complete description of the model is given in **Supplementary Tables S1-4**. Parameter estimation was performed for the six model variants independently and the model fitting against the data is reported in **Figure 1C**. As the number of parameters is different according to the model, we compared their fit quality using the Akaike Information Criterion (AIC) (Akaike, 1974), which penalises models with a higher number of estimated parameters (**Supplementary Table S5**). The two models fitting the data featured wortmannin-regulated ATG13 accumulation and event-triggered disappearance. Since no significant differences were evident in the fitting of either models 1 and 3, we selected model 3 (where wortmannin affects both aggregation and disappearance), as it offered a more general biological mechanism. This model was also the only one where all the parameters were identifiable with a 66% confidence level (**Supplementary Table S3**). Surprisingly, parameter estimation revealed a value of 1.01365 for the parameter *m* (confidence interval at 66%: [0.164715, 1.60609]) representing the partial reaction order of ATG13 complex accumulation on the already aggregated complex. This suggests significant cooperativity for ATG13 (and likely ULK complex) accumulation. Details for parameter estimation and identifiability are reported in the **Materials and Methods** and **Supplementary Table S3.**

**Figure 1.**
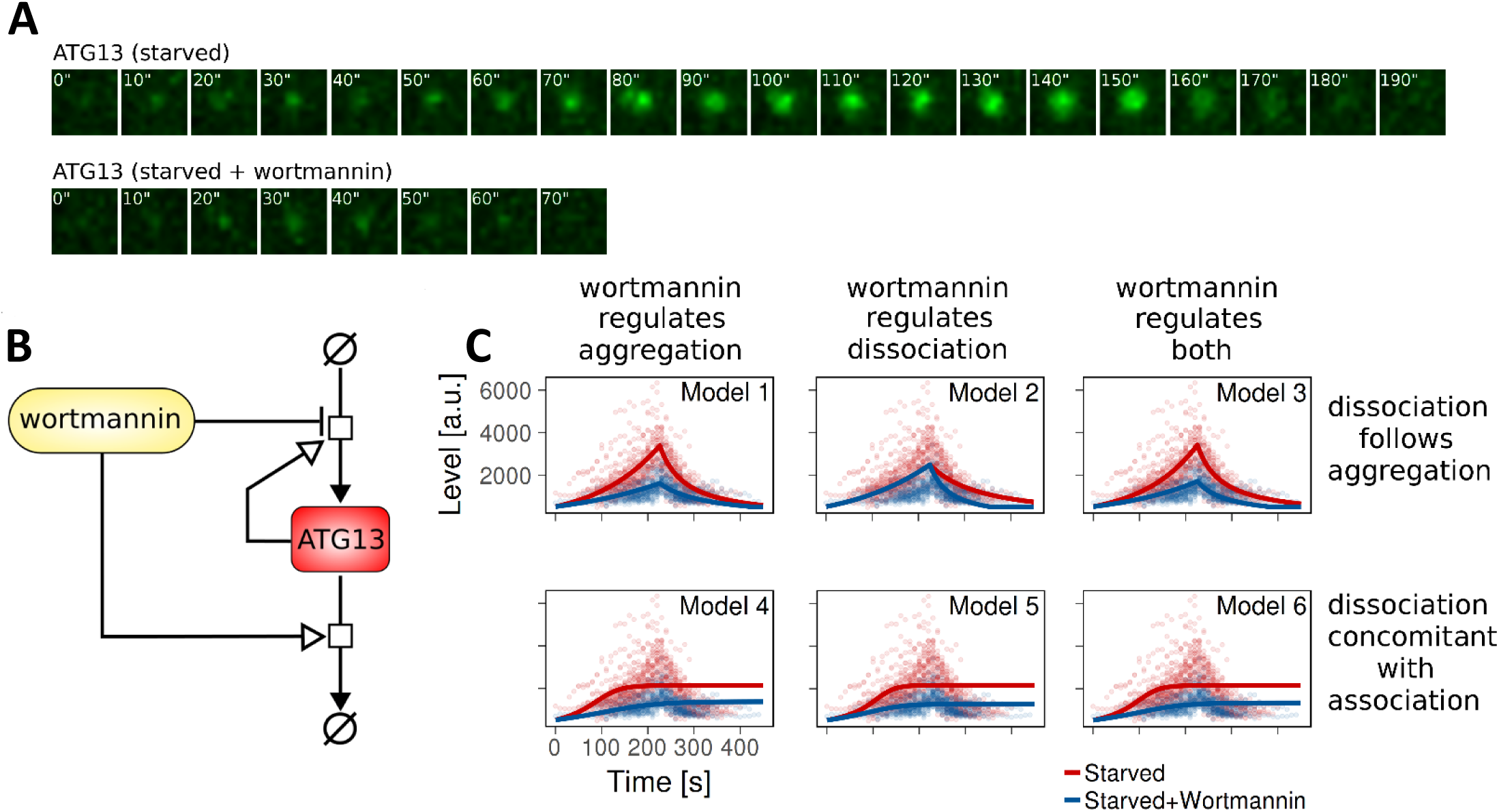
A mathematical model for non-selective autophagy. A. Representative time course images of one autophagy event involving the formation and disappearance of a GFP-ATG13 punctum. In the first row, cells are starved. In the second row, cells are starved and treated with wortmannin. B. Schematic diagram of the mathematical model for non-selective autophagy. Please see text for details. C. Parameter fitting for the 6 model variants. The switch between ATG13 accumulation and removal is regulated by events in Models 1-3. Models 4-6 are eventless. Wortmannin downregulates ATG13 accumulation (Models 1 and 4), ATG13 removal (Models 2 and 5), and both of the reactions (Models 3 and 6). Dots are experimental time points, whereas lines indicate model simulation after parameter estimation. Red indicates starvation, blue indicates starvation plus wortmannin.

### Wortmannin affects ATG13 peak time distribution in non-selective autophagy

Since a PI3P feedback during initiation of autophagosome formation had been suggested, we analysed the effect of wortmannin on ATG13 accumulation (**Figure 2A**,**B**). First, we verified that there was no significant correlation between the signal peak times and initial signal intensities, which could have pointed to a possible misidentification of the start of aggregation and therefore of signal peak times (**Supplementary Figure S2**).The signal intensity of each experimental time course repeat was then normalised within [0,1]. Sorting repeats by decreasing peak times (dark red points in **Figure 2A**,**B**) suggests that the addition of wortmannin changes the probability distribution. Q-Q plots for ATG13 peak times upon starvation (**Figure 2C**) and starvation plus wortmannin (**Figure 2D**) versus normal or log-normal distributions suggested that the peak times for ATG13 fluorescence signal upon starvation tended to approximate a normal distribution, whereas wortmannin shifted the peak times to approximate a log-normal distribution. We statistically tested whether the peak times were normally distributed. The Shapiro-Wilk test could not reject the null hypothesis that the starvation without wortmannin was normally distributed (*p*-value: 0.432; skewness: 0.312, excess kurtosis: −0.251), but did reject the null hypothesis for the starvation plus wortmannin (*p*-value: 0.004; skewness: 1.193, excess kurtosis: 1.716). If the data are log-normally distributed the logarithms of the data are expected to be normal. The Shapiro-Wilk test using the logarithms of the starvation plus wortmannin group could not reject the null-hypothesis (*p*-value: 0.855), suggesting that this data set could be normally distributed. The skewness and excess kurtosis for this set were very close to 0 (skewness: −0.116, excess kurtosis: - 0.198), indicating that the distribution is almost normal and supporting the original conclusion that the starvation plus wortmannin group is log-normally distributed. Using model 3 described above, we simulated populations of non-selective autophagy events with and without wortmannin, where the peak time (parameter *t*) was sampled from a normal or log-normal distribution respectively (**Figure 2E, Supplementary Table S6**). Those simulations adequately reproduce the experimental data and suggest that our model can be used to describe the ATG13-dependent initial step of autophagosome formation.

**Figure 2.**
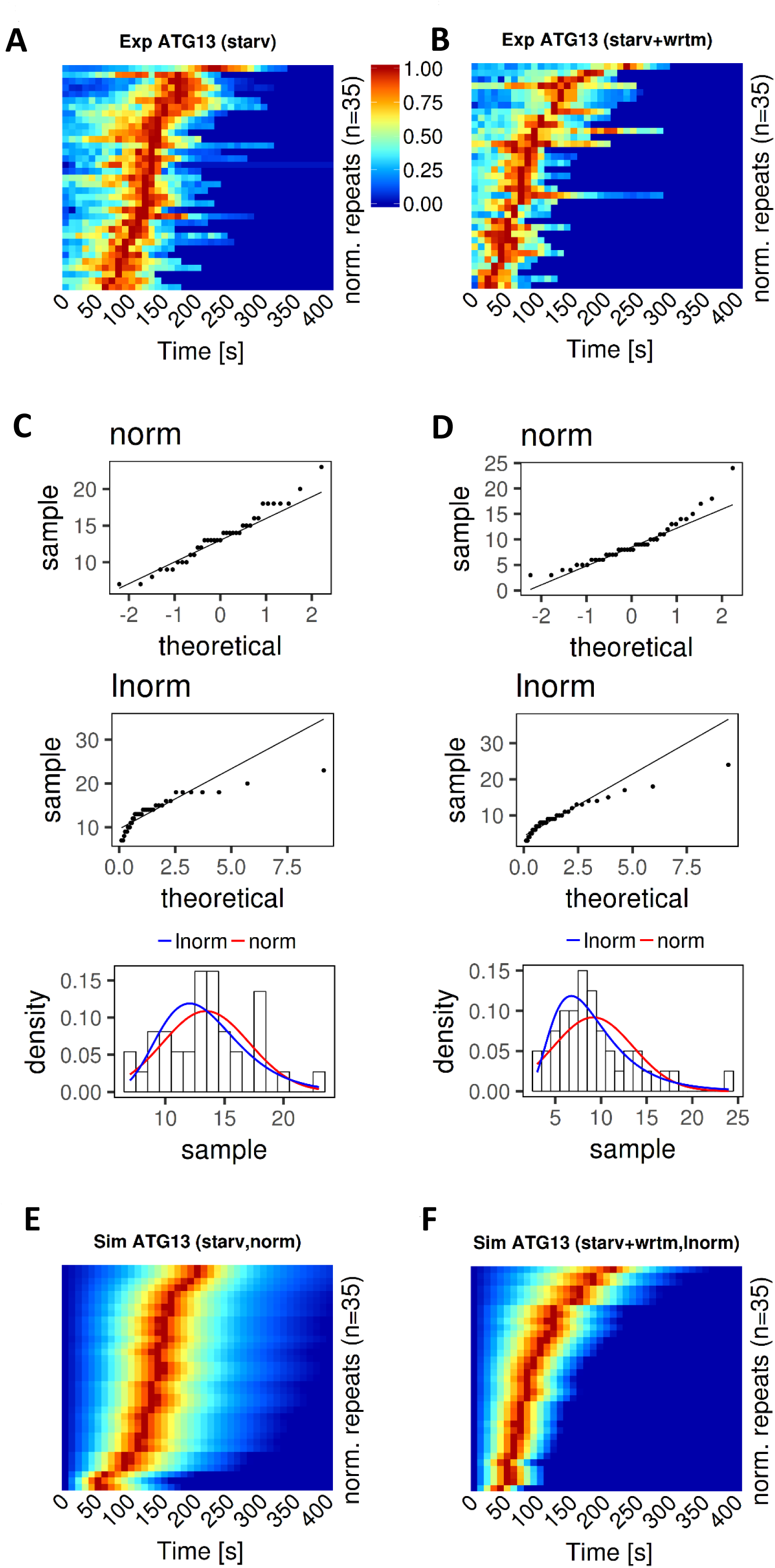
PI3P inhibition upon Wortmannin shifts ATG13 peak time distribution from normal to log-normal. A. Experimental kymographs of ATG13 upon starvation. For each group, n=35 experimental repeats were selected and the signal intensity of each repeat was normalised within [0,1] and indicated with the colours [blue, red]. Repeats were sorted by decreasing peak time. B. Equivalent to panel A but for ATG13 upon starvation plus wortmannin. C. Normality and log-normality analysis of experimental peak times for ATG13 time courses upon starvation confirmed that the peak times were normally distributed. D. Equivalent to panel C but for ATG13 upon starvation plus wortmannin. Peak times for ATG13 were log-normally distributed. E. Simulated kymographs of ATG13 upon starvation. ATG13 peak time (parameter *t*) was sampled from a normal distribution. A total of n=35 simulated repeats were computed and the signal intensity of each repeat was normalised within [0,1] and indicated with the colours [blue, red]. Repeats were sorted by decreasing peak time. F. Equivalent to panel E, but for ATG13 upon starvation plus wortmannin. ATG13 peak time (parameter *t*) was sampled from a log-normal distribution using meanlog and sdlog.

### A mathematical model for mitophagy

To compare and contrast our data from non-selective autophagy to mitophagy we set up live imaging experiments of ATG13 and a mitochondrial marker after treatment with the drug ivermectin (see Zachari et al, 2019). To consider only mitophagy events in our analysis, we ensured that ATG13 co-localised with the targeted mitochondrion and formed, at least for part of the sequence, a quasi-ring structure around the mitochondrion at some point. With these restrictive conditions, we were able to find 23 time courses. Of highest concern to be excluded were events of non-selective autophagosome formation in close apposition to the mitochondria surface (Hailey et al, 2010).

In contrast to the non-selective autophagy where there was a single peak of ATG13, in mitophagy ATG13 appeared to join and leave the targeted mitochondrion several times, revealing oscillatory dynamics (**Fig 3A** shows an example from a live imaging experiment at high frame rate). Time courses were quantified, synchronised, filtered and regularised (see **Materials and Methods** for details). **Figure 3B** shows three such examples and **Figure 3C** shows the final mean time course from the 17 datasets including confidence interval of the mean at 95% confidence level and 1 standard deviation. From the mean time course, we identified the peaks and troughs for each aggregation event and we showed that the delay between events tends to increase exponentially along the time course (**Figure 3D**).

**Figure 3.**
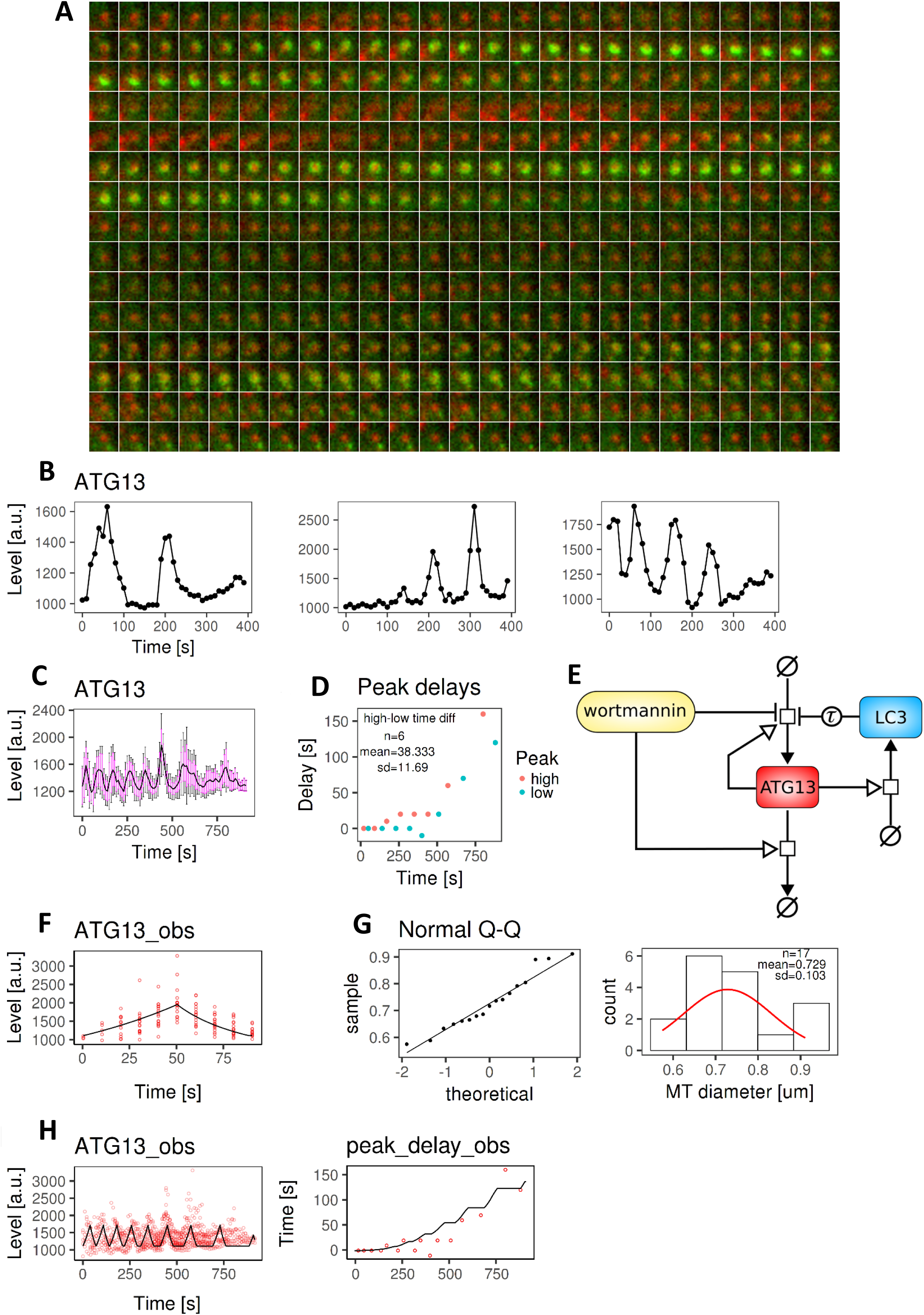
A mathematical model for mitophagy. A. Live imaging of GFP-ATG13 (green) translocations on a targeted mitochondrion (red) showing sequential translocations. Each frame is 0.8’’ and the entire sequence is shown. This is wide field imaging not used for the quantitations but providing a more continuous view of the translocations. B. Three representative regularised quantified time courses up to 400s. These plots are derived from spinning disk confocal images. C. ATG13 mean time course (black line) after data synchronisation, filtering, and regularisation. The mean is shown with the confidence interval of the mean at 95% confidence level (magenta bars) and 1 standard deviation (black bars). D. The delays of the upper and lower aggregation peaks, indicating how these increased over time exponentially. Mean and standard deviation of the time differences between upper and lower peaks are reported as annotations. These statistics define the model parameter *t*. E. Model diagram for mitophagy events. This model extends the diagram in **Figure 1B** by including LC3 and the events for regulating the delays between aggregations. The concentration of LC3 was calculated before the event starting ATG13 accumulation was triggered. Therefore LC3 inhibits ATG13 accumulation after a delay (Tau). F. Model fitting for ATG13 kinetic rate constants using the 1st aggregation data set. The experimental data and model simulation are represented with red circles and black line, respectively. G. MT diameter density and QQ plot against a normal distribution. 17 mitochondrial diameters were measured and analysed. The reported mitochondria diameter mean and standard deviation define the model parameter *MT_diam*. H. Model fitting for ATG13 and the delay between aggregations using the data sets in panel C. The parameters *t* and *MT_diam* were fixed to their mean value during this stage of parameter estimation.

Like non-selective autophagy, the ULK complex initiates a cascading mechanism that leads to mitochondrion engulfment by the LC3-covered membrane. To explain the increasing delays between repeated peaks of ATG13, we hypothesised that ULK complexes can only aggregate in regions of the mitochondrion surface not already covered by LC3 containing membrane. In this view, as engulfment progresses, the portion of mitochondrial surface not covered by LC3 decreases, and the probability of a new ATG13 translocation event decreases. This process would continue until the whole mitochondrion has been engulfed by LC3 containing membrane. We extended the model developed for non-selective autophagy to include LC3 (**Figure 3E**) whose accumulation is driven by the presence of ATG13-containing complexes (see also Karanasios et al, 2013), and it in turn feeds back on those complexes by decreasing the probability of initiating a new aggregation event. This is consistent with the extensive live imaging data we recently reported (Zachari et al, 2019). The complete description of the model is given in **Supplementary Tables S7-10**. This model was fitted with the mitophagy time course data. First, to parametrise ATG13 aggregation and disappearance, we extracted the first peak of each time course and synchronised them as we did for the non-selective autophagy data set (**Figure 3F, Supplementary Table S9**, and **Supplementary Figure S3.** See **Materials and Methods** for details). Parameter estimation revealed that the ATG13 kinetic rate constants were not significantly different between the mitophagy model (parameters: kprodATG13 = 0.0114, CI95 = [0.0099, 0.0127]; kremATG13 = 0.0114, CI95 = [0.0010, 0.0139]) and the non-selective autophagy model (parameters: kprodATG13 = 0.0082, CI95=[0.0052, 0.0193]; kremATG13 = 0.0029, CI95=[0.0007, 0.0183]). We then estimated the remaining model parameters using the complete ATG13 time course data set. In the model, there are two parameters sampled from random variables. The first is the time between the start of an aggregation event and the peak of ATG13 intensity, which is sampled at each aggregation (parameter *t* as in the non-selective autophagy model). In line with non-selective autophagy (**Figure 2C**), we assumed a normal distribution for ATG13 peak times. We directly extracted the information needed to compute *t* from the time differences between the upper and lower peaks in **Figure 3D**. Detailed statistics are reported in **Supplementary Table S11**. The second parameter is the mitochondrial diameter which is randomly sampled at the beginning of each time course from the distribution of measured mitochondrial diameters. By analysing the mitochondrial diameters from the imaging time courses, we found that the population approximated a normal distribution (**Figure 3G, Supplementary Figure S4**). In agreement with this observation, the Shapiro-Wilk statistical test could not reject the null hypothesis that these data were normally distributed (*p*-value: 0.379; skewness: 0.378, excess kurtosis: - 1.061). In order to fit the mean experimental time course (**Figure 3C**) we fixed the values of time to peak and mitochondrial diameter to the mean of their experimental distributions as reported in **Figures 3D, 3G (second plot). Figure 3H** shows the fitting between model and data. Details for parameter estimation and identifiability are reported in the **Materials and Methods**. Parameter estimation gave an insight into the increasing delay between aggregations. In particular, the parameter *p* determining the exponential increase for the delay was estimated to be ∼2.79. This cubic growth is consistent with the idea that each aggregation event represents coating of the target by both ATG13 and LC3, and that as more of the area will be covered there will be a delay until the next available region is found and the LC3 modifying machinery translocates there. We tested for this in the next section.

### Mitochondrial fragment diameter determines the number of ATG13 translocations

According to our hypothesis, each occurrence of ATG13 translocation (aggregation) promotes a partial coverage of a mitochondrion by LC3-containing membrane, and this process terminates when the whole mitochondrion becomes engulfed (see **Figure 4A**). According to this, the increasing delay between observed ATG13 peaks is due to two phenomena: 1) the time spent by ATG13 (as part of the ULK complex) searching a region on the mitochondrion surface which has not yet been covered by LC3 membrane, and 2) the translocation of the LC3-modifying machinery. Under the assumption that the ULK complex drives engulfment by LC3, the latter delay should be constant over time as no random search is involved. Therefore, our model explicitly represents only the ATG13-dependent delay.

**Figure 4.**
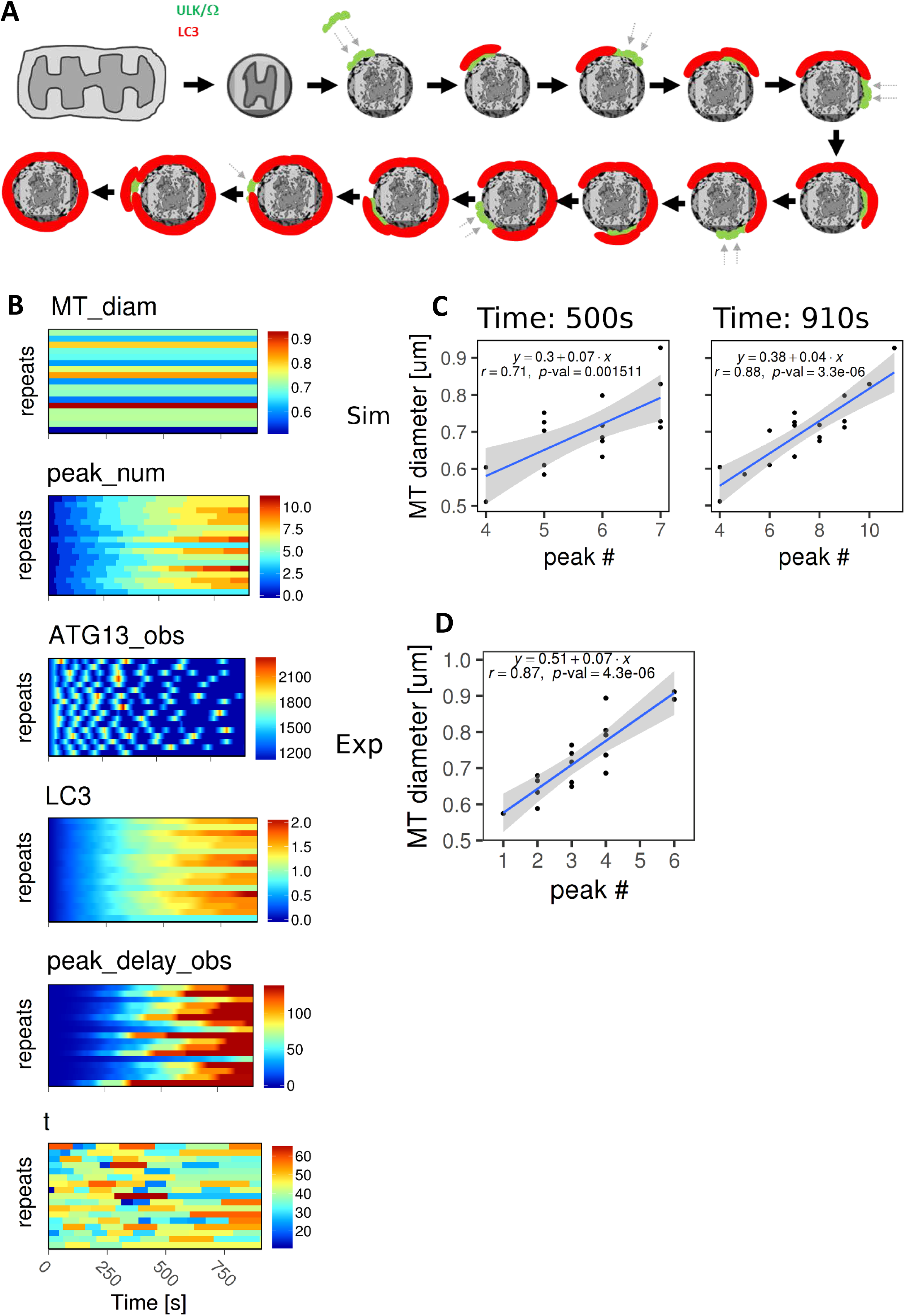
ATG13 sequential translocations depend on the mitochondrial size. A. Proposed model for the sequential translocations of ATG13 on the targeted mitochondrial fragments. After fragmentation and damage, the early autophagic machinery represented by the ULK complex and omegasomes assembles on a piece of the mitochondrion surface (green elements) to be followed by LC3 membrane formation (red). This process is repeated 6 times until the mitochondrial fragment is fully engulfed by the LC3 membrane. Arrows indicate novel translocations to a new mitochondrial area. B. A population of 17 mitophagy events was simulated by sampling the mitochondrial diameter (model parameter: *MT_diam*) and ATG13 accumulation time (model parameter: *t*) from their normal distributions, respectively. The single time course repeats are plotted by row on each plot. The signal intensity is depicted with a coloured scale. ATG13 signal is shown to oscillate in each repeat and these aggregations gradually become unsynchronised due to the accumulation of delay between events. C. The model predicted a linear positive correlation between the mitochondrial diameters and the number of ATG13 aggregations at the end of the simulation (910s). The fitting line is indicated in blue, whereas its confidence interval in grey. D. Experimental relationship of mitochondrial diameter and number of ATG13 aggregations, reporting very similar correlation to the simulation in C.

Using the parameters estimated above, we simulated 17 mitophagy events with the parameter *MT_diam* - determining the mitochondria diameter within the model – sampled from a normal distribution inferred from the data at the beginning of each simulation (see distribution parameters in **Figure 3G plot 2, Supplementary Table S10**). The parameter *t*, setting the time between the beginning and the maximum signal of each ATG13 aggregation event, was also sampled from a normal distribution inferred from the data at the beginning of each ATG13 accumulation (see distribution parameters in **Figure 3D, Supplementary Tables S10-11**). The single simulation repeats for the main model readouts are shown in **Figure 4B**. The oscillations for a population of mitophagy events tend to lose synchronisation over time due to the stochastic increase in delay between peaks. The normal distribution of aggregation duration means that the quantity of LC3-membrane for each aggregation event driven by ULK complex will also be normal. This suggested that the size of a mitochondrion could determine the total number of accumulations needed to fully engulf it. Therefore, we analysed whether there was a correlation between the number of ATG13 peaks and the sampled mitochondrial diameters. This analysis was conducted at the end of the simulation (910 s), and in the middle (500 s). Remarkably, the simulated data predicted a significant linear positive correlation (r = 0.88, *p*-value = 3.3e-0.6) between the number of ATG13 aggregations and the diameters of the corresponding engulfed mitochondria at the end of the simulation (**Figure 4C**). To test this prediction, we counted the number of peaks for the 17 repeats in our experimental data and plotted them against the corresponding mitochondrial diameters. A significant linear positive correlation was again obtained (r = 0.87, *p*-value = 4.3e-06) (**Figure 4D**).

## Discussion

In recent years, we and others have described the early membrane re-arrangements involved in the initiation of autophagy in some detail (Koyama-Honda et al. 2013; Ktistakis and Tooze, 2016; Zhao and Zhang, 2018). It is very clear from that work that the process is extremely complicated, it involves a series of hierarchical translocations of the relevant components on and off the forming autophagosomes, and it uses a large number of proteins and protein complexes. Here, we have focused on one of the earliest steps in this pathway, the translocation of the ATG13 protein to the forming autophagosomal structure, and used mathematical models to gain insight into the mechanisms that regulate it.

Formation of ATG13 puncta in non-selective autophagy is characterised by a single event of accumulation. In our model, the cooperative aggregation of ATG13 is represented by the model parameter *m*, increasing the partial order of the ATG13 species for the reactions of accumulation and disappearance. Model parameter estimation partially identified this parameter and calculated a positive value, supporting the hypothesis of a positive feedback loop increasing and decreasing the rate of ATG13 accumulation and removal, respectively. Such cooperativity could be explained for instance by a surface effect: the larger the ATG13 (and by extension the ULK complex) aggregates, the more sites there are for aggregation or disassembly.

Wortmannin-dependent inhibition of the VPS34 complex, which regulates PI3P production, led to a decrease in duration and intensity of ATG13 aggregations and a change in the probability distribution of ATG13 peak times from normal to log-normal. One possible explanation is that inhibition of PI3P synthesis removes external inputs on ULK complex accumulation, letting the process be driven only by complex stability. Without input from external factors, many natural phenomena relating to frequencies, for example, in this case, the kinetics of ATG13 disappearance (**Figure 2D**), can be described and explained by a log-normal distribution (Hosoda et al, 2011; Limpert et al, 2001).

In contrast to non-selective autophagy, in mitophagy ATG13 accumulation and disappearance were more complex and displayed oscillatory dynamics (**Figure 3A**). According to our hypothesis, each event is characterized by ATG13 aggregation on an LC3-free subsurface on the targeted mitochondrion and drives the subsequent coverage by LC3 membrane of this region (**Figure 4A**). Using time-course data analysis and mathematical modelling, we showed that these events delay exponentially along the time course. In addition, model simulations predicted, and experimental data confirmed, that the number of sequential translocations (oscillations) is proportional to the mitochondrial diameter. These findings indicate that mitophagy is dependent on the target structure and that the ULK complex as exemplified by the ATG13 protein is a key component for this process. How does this complex find an LC3-free subsurface on the mitochondrion? A potential explanation could be that the ULK complex can only accumulate on a region on the mitochondrion where mitophagy receptors can be detected. In this view, a region partially engulfed by LC3 would prevent the binding between the ULK complex and the mitophagy receptors.

Our study also analysed the delay between ATG13 sequential translocations. Model parameter estimation revealed that this delay increases almost cubically with time. Given that the complex covers a surface, the delay between each step of partial engulfment would be expected to grow quadratically if no delays were involved. However, as the number of ATG13 translocations inherently depends on the time it takes the LC3 machinery to partially engulf the targeted mitochondrion, it appears clear that the translocation and engulfment of the LC3 machinery onto the mitochondrion surface adds a delay, and that our model calibration estimates the sum of the ATG13- and LC3-dependent delays. Our future work will attempt to introduce additional components to our imaging experiments to expand the scope (and complexity) of these models.

## Materials and Methods

### Cell cultures

HEK-293 cells stably expressing GFP-ATG13 have been described before (Karanasios et al, 2013). Immunofluorescence microscopy was performed as described (Karanasios et al, 2015). Mitophagy experiments were performed as described (Karanasios et al, 2016, and Zachari et al, 2019).

### Raw data acquisition

Images were acquired using a spinning disk confocal microscope, comprising Nikon Ti-E stand, Nikon 100x 1.49 NA objective, Yokogawa CSU-X scanhead, Andor TuCam splitter and Andor iXon 897 EM-CCD cameras. Using the splitter and 2x cameras enabled GFP (ATG13) and mCherry (mitochondria) images to be captured simultaneously. Images comprising 512×512 160 nm pixels were taken using a 200-300 ms exposure time, with z-stacks acquired at 10 s intervals. Each stack covered ∼6 μm range, with the step interval set at 500-700 nm to ensure optimal sampling. Maximum intensity projections of cropped regions were created using FIJI (Schindelin et al, 2012) and analysed using Imaris software v9.1.2 (http://www.bitplane.com) to quantify mean signal intensities along the time course. To generate the non-selective autophagy data set, raw confocal images were acquired from 8 cells. From these images, 37 non-selective autophagy events were selected upon starvation and 40 events upon starvation plus wortmannin, measuring ATG13 as readout. As autophagy events were triggered within the cytoplasm at different time points, we synchronised the time courses based on their maximum intensity peak. To generate the mitophagy data set, raw confocal images were acquired from 25 cells and 23 events were selected assuring that ATG13 (green channel) co-localised with a mitochondrion (red channel) and ATG13 formed a quasi-circle around the mitochondrion at some point within the time course. This allowed us to exclude potential autophagosome formation on the mitochondrion surface. This selection of mitophagy events was then quantified (**Supplementary Figure S5**). Then, the time courses were manually synchronised, overlapping the delays between aggregations. To facilitate the synchronisation, the time courses were splined reducing the noise within the data (**Supplementary Figure S6**). Once synchronised, the splined time courses were replaced with the original time courses (**Supplementary Figure S7, AB**). Six time courses were removed due to their highly irregular nature. The initial and late time points were also cut off in order to have at least three repeats for each time point (**Supplementary Figure S7, C**). Finally, the time courses were regularised to eliminate the gradual signal decline typically occurring over time in fluorescence images (**Supplementary Figure S7, D**). The final single regularised time courses are reported in **Supplementary Figure S8**.

Mitochondria diameters were extracted using ImageJ 1.51 (Schneider et al, 2012). At least 8 mitochondria diameter measurements were taken for each of the 17 time courses (**Supplementary Figure S4**). The averages of these measurements were then computed and summarised in **Figure 3G**.

### ATG13 non-selective autophagy model

Parameter estimation was performed using Copasi 4.22 (Hoops et al, 2006) and SBpipe 4.20.0 (Dalle Pezze and Le Novère, 2017). Particle Swarm optimisation algorithm as implemented in Copasi was configured as follows: iteration number=1000, and swarm size=100. SBpipe was configured to run 1000 parameter estimates using Copasi within the SGE cluster at the Babraham Institute (UK).

The mathematical description for the non-selective autophagy model including its events and parameters is reported in **Supplementary Tables S1-4**. Synchronised time courses for ATG13 upon starvation and starvation plus wortmannin were simultaneously used as data sets for parameter estimation. The parameters (*kprodATG13, kremATG13, m, kwrtm, t*) were estimated for the models 1-3. The same parameters except *t* were estimated for models 4-6, as these models do not include events. For each of the six model variants, a total of 1000 parameter estimates were independently calculated. The 75% of the best fits (reporting the lowest objective value) were selected for analysis. Profile Likelihood Estimation (PLE) analysis was executed at the confidence levels of 66%, 95%, and 99% (**Supplementary Table S3**). Due to the different number of estimated parameters, the AIC was used for ranking the model variants (**Supplementary Table S5**).

### ATG13 mitophagy model

The mitophagy model extended the non-selective autophagy model with the inclusion of LC3 and oscillatory dynamics with delay for ATG13. The mathematical description for the mitophagy model including its events and parameters is reported in **Supplementary Tables S7-10**. Parameter estimation for the mitophagy model was performed in two steps. Firstly, the first aggregation of ATG13 time courses was extracted and synchronised on the peak of maximum signal intensity as for the non-selective autophagy data. This reduced data set was used for estimating the kinetic rate constants for ATG13 (*kprod_ATG13* and *krem_ATG13*), and therefore compute the slopes of ATG13 signalling independently of the complex oscillatory behaviour (**Supplementary Table S9, Supplementary Figure S3A**). Secondly, the parameters regulating the repeated aggregations (*kprodLC3, kpeak, p*) were estimated in two rounds of parameter estimation and identifiability. The first round revealed that *p* and *kprodLC3* were identifiable, although *kprodLC3* minima lay on a defined plateau. Fixing *kprodLC3*, the interdependent parameter *kpeak* was estimated and identified in a second round of parameter estimation (**Supplementary Table S9, Supplementary Figure S3BC**). The parameters *t* and *MT_diam*, representing the time before reaching an aggregation peak and the mitochondria diameter respectively, were fixed to the mean of their normal distribution during the second stage of parameter estimation.

### Statistics

The statistical and programming language R version 3.4.0 was used to generate statistics and plots. AIC (Akaike, 1974) as computed in SBpipe was used for selecting the non-selective autophagy model which fitted the non-selective autophagy dataset. Shapiro-Wilk test was used for testing normality distribution. A batch correction using R package SVA (Leek et al, 2012) was applied to the intensities of LC3 in **Figure 1A** to make the repeats of the two experiments of comparable intensity levels.

## Author Contributions

The project was designed by NLN and NTK. Mathematical modelling was performed by PDP. Live imaging data were collected by EK and SAW from samples prepared by EK and MM. Images were analyzed by PDP and VK. The manuscript was written by PDP, NLN and NTK with contributions from SAW.

