## Supplementary information for "ATG13 dynamics in non-selective autophagy and mitophagy: insights from live imaging studies and mathematical modelling"

### Supplementary Information
### for

ATG13 dynamics in non-selective autophagy and mitophagy: insights from live imaging studies and mathematical modelling.

Piero Dalle Pezze, Eleftherios Karanasios, Varvara Kandia, Maria Manifava, Simon A. Walker, Nicolas Gambardella Le Novère and Nicholas T. Ktistakis.

A) ATG13 ODE for Models 1 and 4

$$\frac{d\left([\mathrm{ATG13}]\cdot V_{\mathrm{ER}}\right)}{d\,t} = +V_{\mathrm{ER}}\cdot\left(\frac{\mathrm{kprodATG13}\cdot[\mathrm{ATG13}]^{m}}{1+\left(\frac{\mathrm{wrtm}}{\mathrm{kwrtm}}\right)^{n}}\right) - V_{\mathrm{ER}}\cdot\left(\mathrm{kremATG13}\cdot[\mathrm{ATG13}]^{1+m}\right)$$

B) ATG13 ODE for Models 2 and 5

$$\frac{d\left([\mathrm{ATG13}]\cdot V_{\mathrm{ER}}\right)}{d\,t} = +V_{\mathrm{ER}}\cdot\left(\mathrm{kprodATG13}\cdot[\mathrm{ATG13}]^{m}\right) - V_{\mathrm{ER}}\cdot\left(\mathrm{kremATG13}\cdot[\mathrm{ATG13}]^{1+m}\cdot\left(1+\left(\frac{\mathrm{wrtm}}{\mathrm{kwrtm}}\right)^{n}\right)\right)$$

C) ATG13 ODE for Models 3 and 6

$$\frac{d\left([\mathrm{ATG13}]\cdot V_{\mathrm{ER}}\right)}{d\,t} = +V_{\mathrm{ER}}\cdot\left(\frac{\mathrm{kprodATG13}\cdot[\mathrm{ATG13}]^{m}}{1+\left(\frac{\mathrm{wrtm}}{\mathrm{kwrtm}}\right)^{n}}\right) - V_{\mathrm{ER}}\cdot\left(\mathrm{kremATG13}\cdot[\mathrm{ATG13}]^{1+m}\cdot\left(1+\left(\frac{\mathrm{wrtm}}{\mathrm{kwrtm}}\right)^{n}\right)\right)$$

D) Assignments for Models 1-6

$$\mathrm{ATG13\_obs} = \mathrm{ATG13\_sf}\cdot[\mathrm{ATG13}]$$

$$\mathrm{wrtm} = \begin{cases} \mathrm{wrtm\_flag} = 0, & \frac{0}{\mathrm{wrtm\_sf}} \\ \mathrm{else}, & \frac{\mathrm{wrtm\_si}}{\mathrm{wrtm\_sf}} \end{cases}$$

$$\mathrm{wrtm\_scaled} = \mathrm{wrtm\_sf}\cdot\mathrm{wrtm}$$

**Supplementary Table S1. ODE table for non-selective autophagy models 1-6.**

The ODE regulating ATG13 dynamics differs among model 1-6, depending on the positive interaction between PI3P and ATG13. As the model does not include PI3P explicitly, PI3P inhibitor wortmannin was modelled to directly inhibit ATG13. Models 1-3 and 4-6 share the same ODE, correspondingly, but models 4-6 do not include events. A) Wortmannin downregulates ATG13 accumulation. B) Wortmannin upregulates ATG13 removal. C) Wortmannin downregulates ATG13 accumulation and upregulates ATG13 removal. D) The assignment variables shared in all the model variants. *ATG13_obs* is the only model observable which is associated with the experimental data during parameter estimation. *Wrtm_flag* is a boolean variable indicating the presence or absence of wortmannin in the model. When set to 1, *wrtm* negatively affects ATG13 dynamics.

**A) ATG13_accumulation (fire at start)**

Trigger expression

Time < t

Target

kprodATG13 ← kprodATG13.InitialValue

kremATG13 ← 0

**B) ATG13_removal**

Trigger expression

kprodATG13 > 0 and kremATG13 = 0

Delay (calculation and assignment)

t

Target

kprodATG13 ← 0

kremATG13 ← kremATG13.InitialValue

**C) ATG13_basal**

Trigger expression

kprodATG13 = 0 and kremATG13 > 0 and [ATG13] < 1

Target

kprodATG13 ← 0

kremATG13 ← 0

**Supplementary Table S2. Events table for non-selective autophagy models 1-3.**

Models 1-3 include 3 events which regulate the switch between ATG13 accumulation and removal. A) The estimated parameter *t* represents the time point when the switch between the two reactions happens. While *Time* is lesser than *t*, ATG13 can only accumulate. B) ATG13 removal is regulated by a delay event which is triggered after *t* seconds of ATG13 accumulation. C) Once ATG13 is below a certain level (after ATG13 removal), ATG13 returns to its basal level.

| Model | Parameter | Value | LeftCI66 | RightCI66 | LeftCI95 | RightCI95 | LeftCI99 | RightCI99 |
|---|---|---|---|---|---|---|---|---|
| m1 | kprodATG13 | 0.00932224 | 0.0062057 | 0.0193744 | 0.00534413 | 0.0204578 | 0.00485426 | 0.0215034 |
| | kremATG13 | 0.00364467 | 0.00130477 | 0.0131757 | 0.000925171 | 0.0200528 | 0.000669717 | 0.0263238 |
| | kwrtm | 1.36189 | 0.633578 | 2.53802 | 0.448353 | 2.95053 | 0.404588 | 3.58819 |
| | m | 0.872221 | -Inf | 1.37514 | -Inf | 1.60431 | -Inf | 1.7034 |
| | t | 225.965 | 210.457 | 241.2 | 203.19 | 250.158 | 193.289 | 257.067 |
| m2 | kprodATG13 | 0.00815308 | 0.00408281 | 0.0161227 | -Inf | Inf | -Inf | Inf |
| | kremATG13 | 0.00261136 | 1.92E-05 | 0.0131946 | -Inf | 0.0223888 | -Inf | 0.0381103 |
| | kwrtm | 0.603469 | 0.0266956 | Inf | -Inf | Inf | -Inf | Inf |
| | m | 0.79801 | -Inf | 2.40739 | -Inf | Inf | -Inf | Inf |
| | t | 224.952 | 150.113 | 246.585 | -Inf | 259.689 | -Inf | 269.753 |
| m3 | kprodATG13 | 0.00822279 | 0.00528378 | 0.0168838 | 0.00523032 | 0.0192829 | 0.00484189 | 0.0202791 |
| | kremATG13 | 0.00284586 | 0.000990108 | 0.00984171 | 0.000703241 | 0.0182583 | 0.000577289 | 0.0254441 |
| | kwrtm | 1.73235 | 0.860786 | 2.94654 | 0.638663 | 3.97607 | 0.475282 | 4.55836 |
| | m | 1.01365 | 0.164715 | 1.60609 | -Inf | 1.64505 | -Inf | 1.7628 |
| | t | 225.454 | 211.687 | 240.292 | 199.711 | 249.436 | 195.022 | 257.942 |
| m4 | kprodATG13 | 0.0126804 | 0.00493445 | 0.212659 | -Inf | Inf | -Inf | Inf |
| | kremATG13 | 0.00311011 | 0.00126824 | 0.0555853 | -Inf | 2966.03 | -Inf | 3104.35 |
| | kwrtm | 1.8235 | 1.04228 | 2.86918 | 0.836667 | 3.57971 | 0.701413 | 6.65874 |
| | m | 1.85612 | -Inf | 4.39006 | -Inf | Inf | -Inf | Inf |
| m5 | kprodATG13 | 0.00930788 | 0.00473933 | 0.133868 | -Inf | Inf | -Inf | Inf |
| | kremATG13 | 0.00225599 | 0.00122831 | 0.0347083 | -Inf | 2902.18 | -Inf | 3094.07 |
| | kwrtm | 1.47676 | 0.910535 | 2.37755 | 0.75612 | 3.50593 | 0.622059 | 5.9251 |
| | m | 2.2431 | -Inf | 4.23665 | -Inf | Inf | -Inf | Inf |
| m6 | kprodATG13 | 0.0106184 | -Inf | 0.150482 | -Inf | Inf | -Inf | Inf |
| | kremATG13 | 0.00258789 | -Inf | 0.039112 | -Inf | 2870.2 | -Inf | 3002.41 |
| | kwrtm | 3.71053 | 2.45462 | 6.07959 | 1.9224 | 8.95748 | 1.61016 | 11.6276 |
| | m | 2.08336 | -Inf | 5.17864 | -Inf | Inf | -Inf | Inf |

**Supplementary Table S3. Table of estimated parameters for non-selective autophagy models 1-6.**

For each model variant, the estimated parameter values and confidence intervals at confidence levels of 66%, 95%, and 99% are reported. The parameter *m* indicates the degree of ATG13 protein cooperativity during ATG13 accumulation and removal. The kinetic rate constants *kprodATG13* and *kremATG13* are the parameters regulating the mass action reactions for ATG13 accumulation and removal. The parameter *kwrtm* is the kinetic rate constant for wortmannin. The parameter *t* is the time point when the switch between ATG13 accumulation and removal occurs. Model 3 was the only one whose parameters were identifiable within a confidence level of 66%.

| Other Parameters | Value |
|---|---|
| wrtm_flag | 0 or 1 |
| ATG13_sf | 525 |
| ATG13_ini | 525 |
| n | 1 |
| wrtm_ini | 1500 |
| wrtm_sf | 1500 |

**Supplementary Table S4. Additional parameters for non-selective autophagy models 1-6.**

These remaining parameters were not estimated. *Wrtm_flag* is a boolean flag indicating the presence or absence of wortmannin in the model. *ATG13_sf* and *ATG13_ini* are the scaling factor and initial value for ATG13. Their value was set to ATG13 basal level within the data. The parameter *n* indicates the degree of cooperativity for wortmannin. The value of 1 indicates no cooperativity. Finally, *wrtm_ini* and *wrtm_sf* are the scaling factor and initial value for wortmannin. These are currently fixed to a value approximating the average of ATG13 peak intensity upon wortmannin. These values do not affect the model.

| Model | AIC |
|---|---|
| m1 | 5.69E+08 |
| m2 | 8.92E+08 |
| m3 | 5.85E+08 |
| m4 | 8.95E+08 |
| m5 | 8.88E+08 |
| m6 | 8.90E+08 |

**Supplementary Table S5. AIC scores for models 1-6.**

Models 1 and 3 reported the best fitting quality and were the only one fitting the data correctly. Model 3 was chosen as it offered a more general biological mechanism of wortmannin interaction with ATG13.

| Peak times | starvation | starvation+wortmannin |
|---|---|---|
| mean | 134.3243243 | 91.5 |
| sd | 36.70759387 | 43.41511021 |
| skewness | 0.3123986084 | 1.193015561 |
| kurtosis (excess) | -0.2506456494 | 1.715914161 |
| mean * | 4.862389097 | 4.413028409 |
| sd * | 0.2839778206 | 0.4647363216 |
| skewness * | -0.3430226437 | -0.1160637561 |
| kurtosis (excess) * | -0.4434826787 | -0.1979578255 |
| meanlog (for lnorm distrib) ** | 4.864245879 | 4.414814168 |
| sdlog (for lnorm distrib) ** | 0.268370368 | 0.4506102617 |
| * calculated using log(data). | | |
| ** parameters used when sampling from a log-normal distribution. | | |

**Supplementary Table S6. Descriptive statistics for non-selective autophagy data set.**

Descriptive statistics were generated for the time courses. Statistics for the time courses peak times were collected. Skewness and excess kurtosis for the starvation+wortmannin data set indicate how this distribution is not close to a normal distribution compared to the starvation data set, where the values are closer to 0. The same statistics were calculated using the log of the two data sets. Whilst for starvation data set this result indicates a marginal drift from a normal distribution, for the starvation+wortmannin data set the skewness and kurtosis report a closer overlap with a normal distribution, supporting the fact that the original data set was log-normally distributed. We also computed the meanlog and sdlog, which are the parameters for a log-normal distribution from the corresponding normal distribution. The parameters for peak times and lnorm peak times are highlighted and were used for sampling the peak times from a normal or log-normal distribution, respectively, for the simulations reported in Figure 2E-F. Meanlog and sdlog were computed as follows:

*meanlog <- function(mu, v) { log((mu^2)/sqrt(v+mu^2)) }*

*sdlog <- function(mu, v) { sqrt(log(v/(mu^2)+1)) }*

Where *mu* and *v* are the usual mean and variance. Therefore, the log-normal distribution was generated as:

*lognormal* <- *e^normal(meanlog, sdlog)*

$$\frac{d\left([ATG13]\cdot V_{ER}\right)}{dt} = +V_{ER}\cdot\left(\frac{kprodATG13\cdot[ATG13]^{m}}{1+\left(\frac{wrtm}{kwrtm}\right)^{n}}\right) - V_{ER}\cdot\left(kremATG13\cdot[ATG13]^{1+m}\cdot\left(1+\left(\frac{wrtm}{kwrtm}\right)^{n}\right)\right)$$

$$\frac{d\left([LC3]\cdot V_{ER}\right)}{dt} = +V_{ER}\cdot\left(kprodLC3\cdot\frac{[ATG13]}{EC50ATG13+[ATG13]}\right)$$

$$ATG13\_obs = ATG13\_sf\cdot[ATG13]$$

$$wrtm = \begin{cases} wrtm\_flag = 0, & \frac{0}{wrtm\_sf} \\ else, & \frac{wrtm\_si}{wrtm\_sf} \end{cases}$$

$$ATG13\_sf = ATG13\_min$$

$$wrtm\_scaled = wrtm\_sf\cdot wrtm$$

$$EC50ATG13 = \frac{ATG13\_max - ATG13\_min}{2}$$

$$cf = \frac{[LC3]}{MT\_surf}$$

$$peak\_delay\_obs = kpeak\cdot cf^{p}$$

**Supplementary Table S7. ODE table for the mitophagy model.**

This model extends the third model for non-selective autophagy with the inclusion of LC3 and the peak delay observable (variable *peak_delay_obs*). LC3 is positively regulated by ATG13 and its production is arrested by an event triggered when the simulated mitochondrion is engulfed. The peaks delay is calculated from the fraction of mitochondrion surface (parameter *MT_surf=pi*MT_diam^2*) that is covered by LC3 (parameter *cf*). In absence of LC3, *cf* is 0 and there is no delay between peaks. When the mitochondrion is completely engulfed, *cf* is 1 and the delay will be maximised (=*kpeak*). At this stage an event triggers the end of the simulation of ATG13 and LC3.

**A) ATG13_accumulation (fire at start)**
    Trigger expression
        ATG13_obs <= ATG13_min and kremATG13 = 0 and cf < 1
    Delay (calculation and assignment)
        peak_delay_obs
    Target
        kprodATG13 ← kprodATG13.InitialValue
        kremATG13 ← 0
        t ← normal(38.33333333,11.6904519445)
        time_fire ← Time
        kprodLC3 ← kprodLC3.InitialValue

**B) ATG13_removal (fire at start)**
    Trigger expression
        kprodATG13 > 0 and Time >= time_fire + t
    Target
        kprodATG13 ← 0
        kremATG13 ← kremATG13.InitialValue
        peak_num ← peak_num + 1

**C) ATG13_basal (fire at start)**
    Trigger expression
        ATG13_obs <= ATG13_min and kremATG13 > 0
    Target
        kprodATG13 ← 0
        kremATG13 ← 0
        kprodLC3 ← 0
        ATG13 ← ATG13_min / ATG13_sf

**D) MT_engulfed (fire at start)**
    Trigger expression

| | |
|---|---|
| Target | cf >= 1 |
| | kprodATG13 ← 0 |
| | kremATG13 ← kremATG13.InitialValue |
| | kprodLC3 ← 0 |

**Supplementary Table S8. Events table for the mitophagy model.**

ATG13 accumulation and removal events regulate the ATG13 process. The first event is delayed depending on the stage of mitochondrion engulfment by LC3 (*peak_delay_obs*). When triggered, the parameter *t* and *time_fire* which regulate the triggering for ATG13 removal event are also updated. In the two steps of parameter estimations, the parameter *t* is fixed to 50 (parameter estimation for the 1st peak) and 38.33333333 (the mean time difference between upper and lower peaks). The last two events control the end of the simulation when the simulated mitochondrion has been engulfed (*cf* >= 1).

| Step | Round | Parameter | Value | LeftCI66 | RightCI66 | LeftCI95 | RightCI95 | LeftCI99 | RightCI99 |
|---|---|---|---|---|---|---|---|---|---|
| 1st peak | 1 | kprodATG13 | 0.0113491 | 0.0105062 | 0.0122197 | 0.00989427 | 0.0127426 | 0.00950567 | 0.0130256 |
| | | kremATG13 | 0.0113493 | 0.0105583 | 0.0125588 | 0.0099646 | 0.0139121 | 0.0095942 | 0.0148284 |
| n peaks | 1 | kprodLC3 | 0.779693 | 0.315287 | 0.867601 | 0.251537 | 0.999483 | 0.188636 | 1.02112 |
| | | p | 2.78977 | 2.26954 | 3.26076 | 1.91662 | 3.5024 | 1.63998 | 9.25167 |
| | | kpeak | 135.796 | 101.526 | Inf | 72.9858 | Inf | 66.9902 | Inf |
| | 2 | kpeak | 136.137 | 127.481 | 143.216 | 119.836 | 151.299 | 115.046 | 156.433 |

**Supplementary Table S9. Tables of estimated parameters for the mitophagy model.**

Parameter estimation for the mitophagy model was performed in two steps. In the first step, the kinetic rate constants for ATG13 accumulation and removal reactions were estimated using only the first peak time course data. These parameters were identifiable. In the second step, the kinetic rate constant regulating LC3 production and the two parameters regulating the peak delay observable (*p* and *kpeak*) were estimated. *kpeak* was practically non identifiable at this stage because partially related to *kprodLC3*. The identifiable parameters *kprodLC3* and *p* were fixed, whilst *kpeak* was re-estimated (round 2). This allowed us to calculate confidence intervals for all the model parameters.

**A Mitophagy (first step of parameter estimation)**

| Other Parameters | Value |
|---|---|
| wrtm_flag | 0 |
| ATG13_min | 1111.71 |
| t | 50 |
| ATG13_max | 1779.31 |
| time_fire | 0 |
| peak_num | 0 |
| MT_surf | Pi * MT_diam^2 |
| MT_diam | 0.7285371148 |

**B Mitophagy (second step of parameter estimation)**

| Other Parameters | Value |
|---|---|
| t | 38.33333333 |

**C Mitophagy (simulation)**

| Other Parameters | Value |
|---|---|
| t (at each evt) | Normal(38.33333333,11.6904519445) |
| MT_diam | Normal(0.7285371148,0.103021783) |

**Supplementary Table S10. Additional parameters for the mitophagy model.**

The mitophagy model includes other parameters which were not estimated but determined by experimental data analysis. A) Values for the additional parameters during the first step of parameter estimation. The parameter *t* is set to 50 as this is the time when the time courses for the first peak were synchronised. It corresponds to the time when ATG13 accumulation reaction is interrupted and ATG13 is activated. The parameter *MT_diam* is set to the mean of the mitochondria diameters distribution. B) For the second step, the parameter *t* is updated to the mean value of its distribution. C) After parameter estimation, the parameters *t* and *MT_diam* are sampled from their normal distributions. The mitochondrial diameter is sampled at the beginning of each simulation, whereas the parameter t is sampled at each ATG13 aggregation.

| name | value |
|---|---|
| mean | 38.33333333 |
| sd | 11.69045194 |
| skewness | 0.8808986197 |
| kurtosis (excess) | -0.90427325 |
| meanlog (lnorm) | 3.601854141 |
| sdlog (lnorm) | 0.298213678 |

**Supplementary Table S11. Descriptive statistics for the peak times across peaks.**

Descriptive statistics were generated for the time differences between the upper and lower peaks of each peak using the mean ATG13 time course. Despite the peak times for a single peak tend to be normally distributed, the peak times across peaks tend to be less than normal as the skewness and excess kurtosis indicate. We also computed the meanlog and sdlog for the log-normal distribution. The log-normal distribution was computed as described in **Supplementary Table S6**.

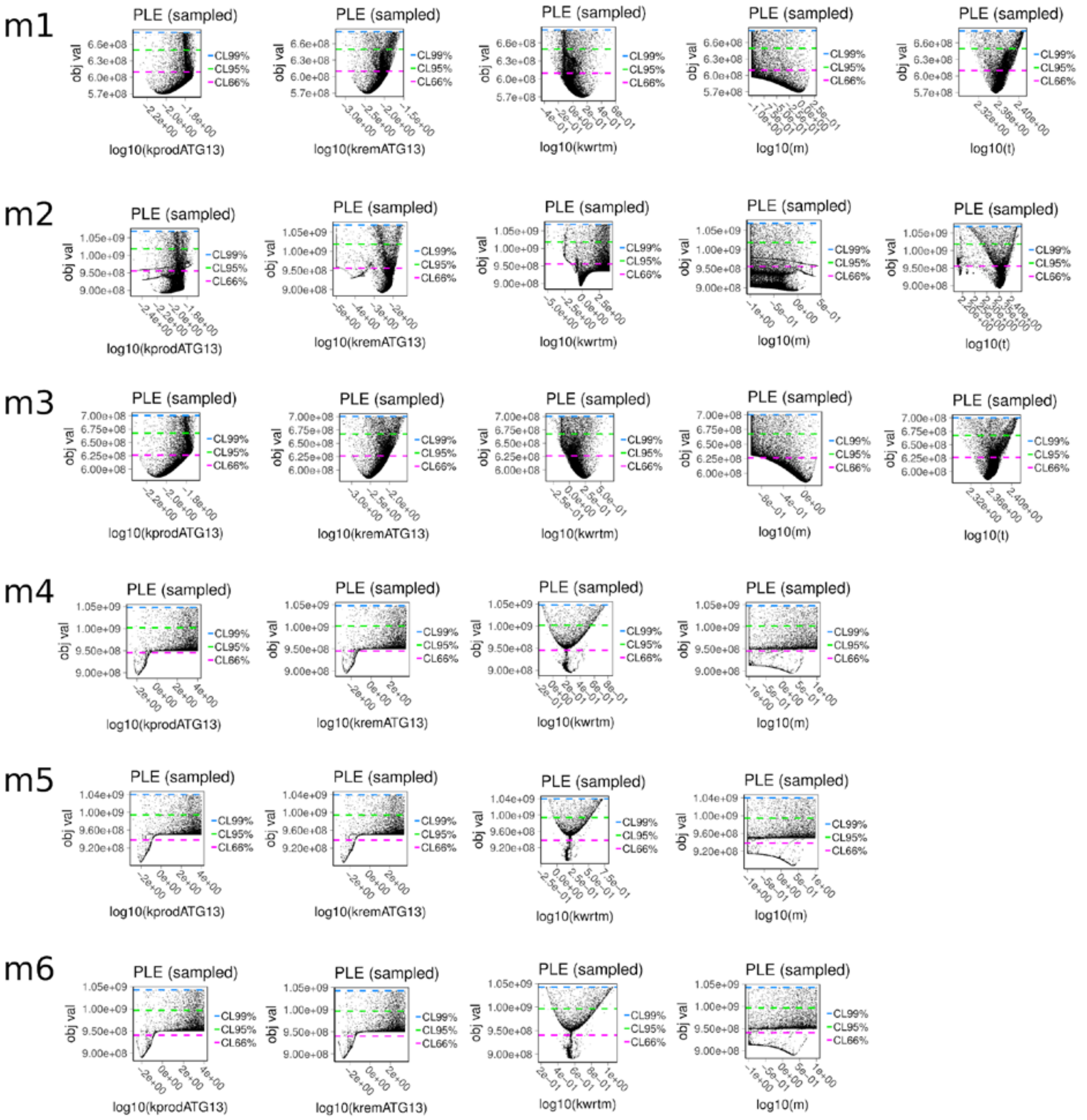

**Supplementary Figure S1. PLE for the parameters of the non-selective autophagy models.**

The PLE plots are represented for each estimated parameters (rows) for the six model variants (columns). For each plot, the x axis reports the parameter values in log10 space, whereas the y axis reports the model objective value. The confidence levels at 66%, 95%, and 99% are reported as coloured horizontal dashed lines. Models 1 and 3 were the only ones to fit the data and model 3 was the only one where each parameter was identifiable within a confidence level of 66%. Model 4-6 did not fit the data and their parameters resulted largely non-identifiable.

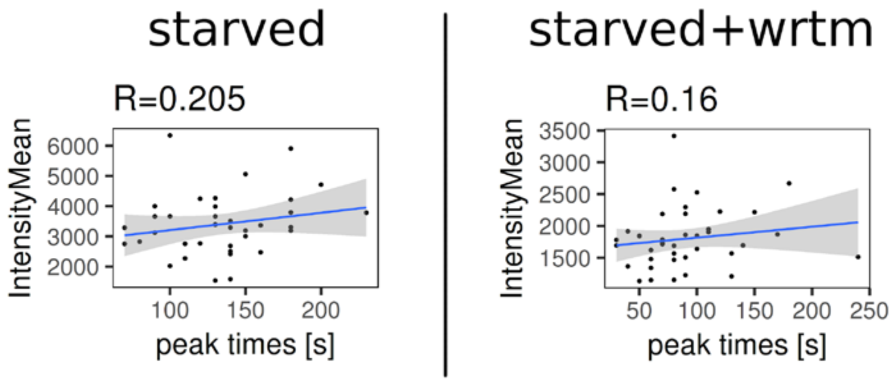

**Supplementary Figure S2. Peak times did not correlate with signal intensities for the non-selective autophagy data.**

No significant correlation was found between peak times and corresponding signal intensities for the starvation and starvation+wortmannin data sets in the non-selective autophagy.

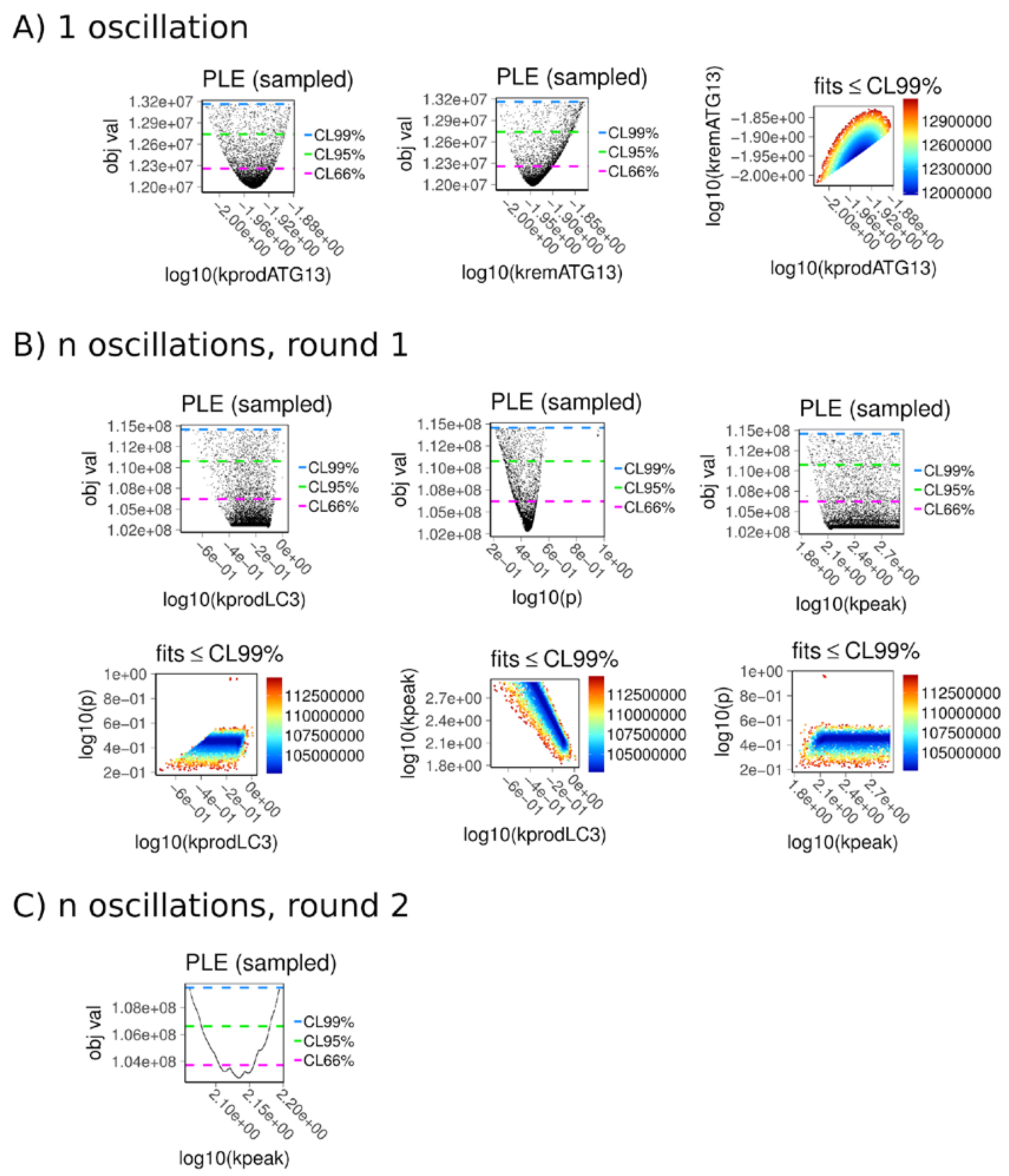

**Supplementary Figure S3. PLE for the parameters of the mitophagy model.**

A) Parameter estimation and identifiability using the first peak data set. ATG13 kinetic rate constants were identifiable within a confidence level of 99% and did not correlate with each other.

B) First round of parameter estimation and identifiability using the complete time course data set. In the first round of parameter estimation, the parameters *p* and *kprodLC3* were identifiable although *kprodLC3* minima lay on a plateau. The parameter *kpeak* was practically non-identifiable (see infinite upper bound).

C) After fixing the parameters *p* and *kprodLC3*, the parameter *kpeak* was re-estimated and found identifiable.

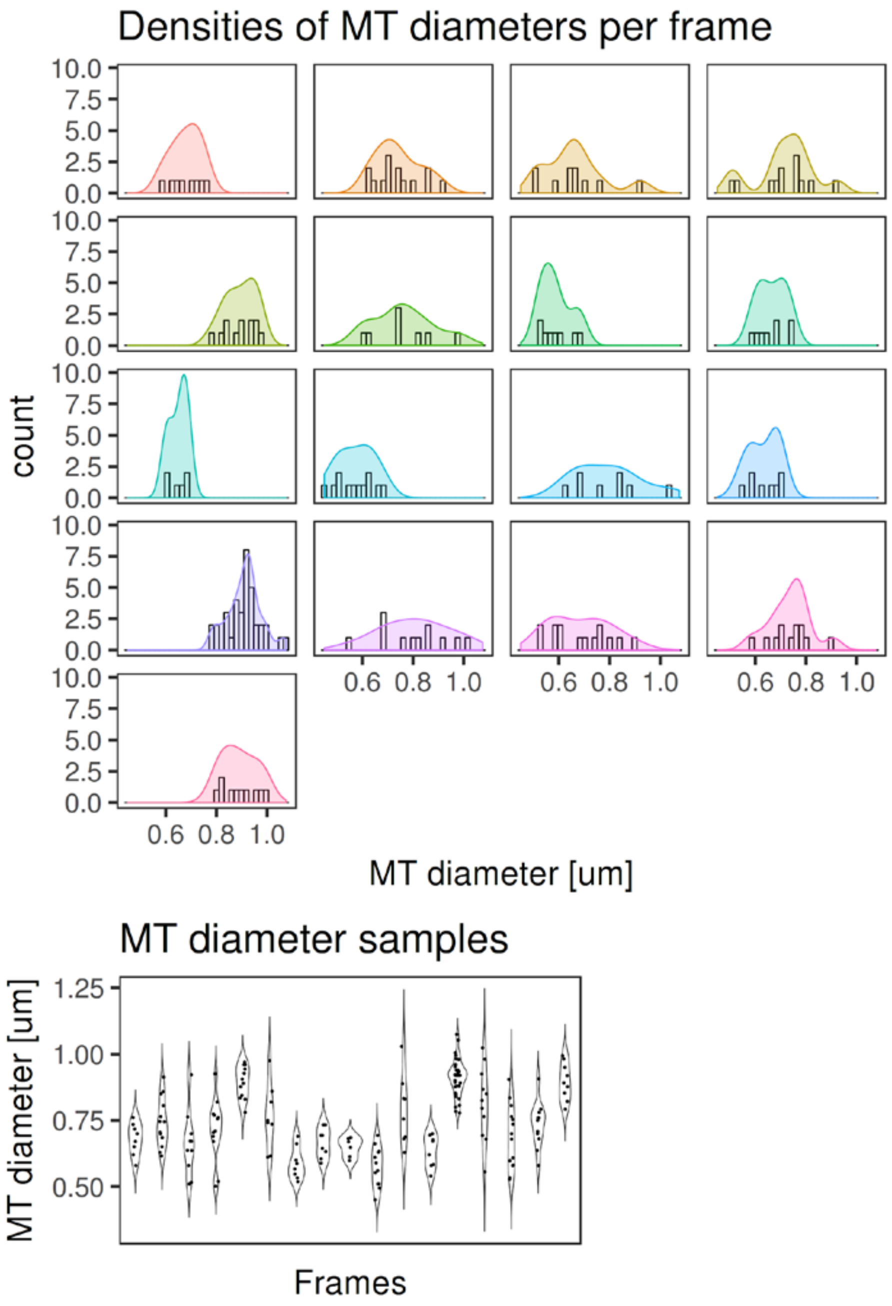

**Supplementary Figure S4. Mitochondrial diameter measurements for each frame.**

The diameter of the mitochondria engulfed by ATG13 was measured for each of the 17 time courses. The mean diameter for each frame was then computed.

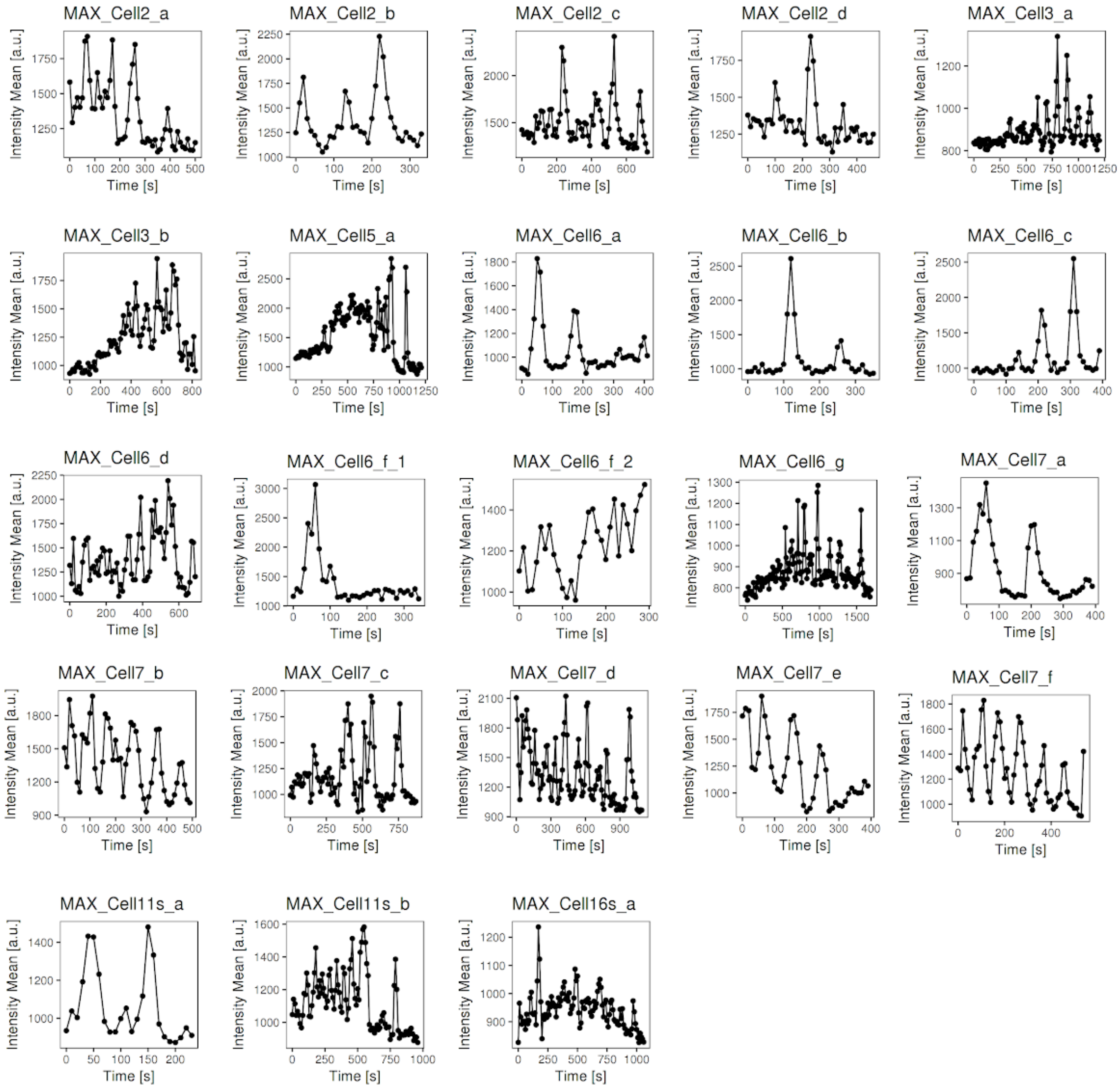

**Supplementary Figure S5. Raw quantified time courses for ATG13 for the mitophagy model.**

A total of 23 time courses were quantified from fluorescence large images. ATG13 accumulations/removals have similar lengths but the time between two peaks tend to increase over time. Due to this regularity, the time courses could have been synchronised.

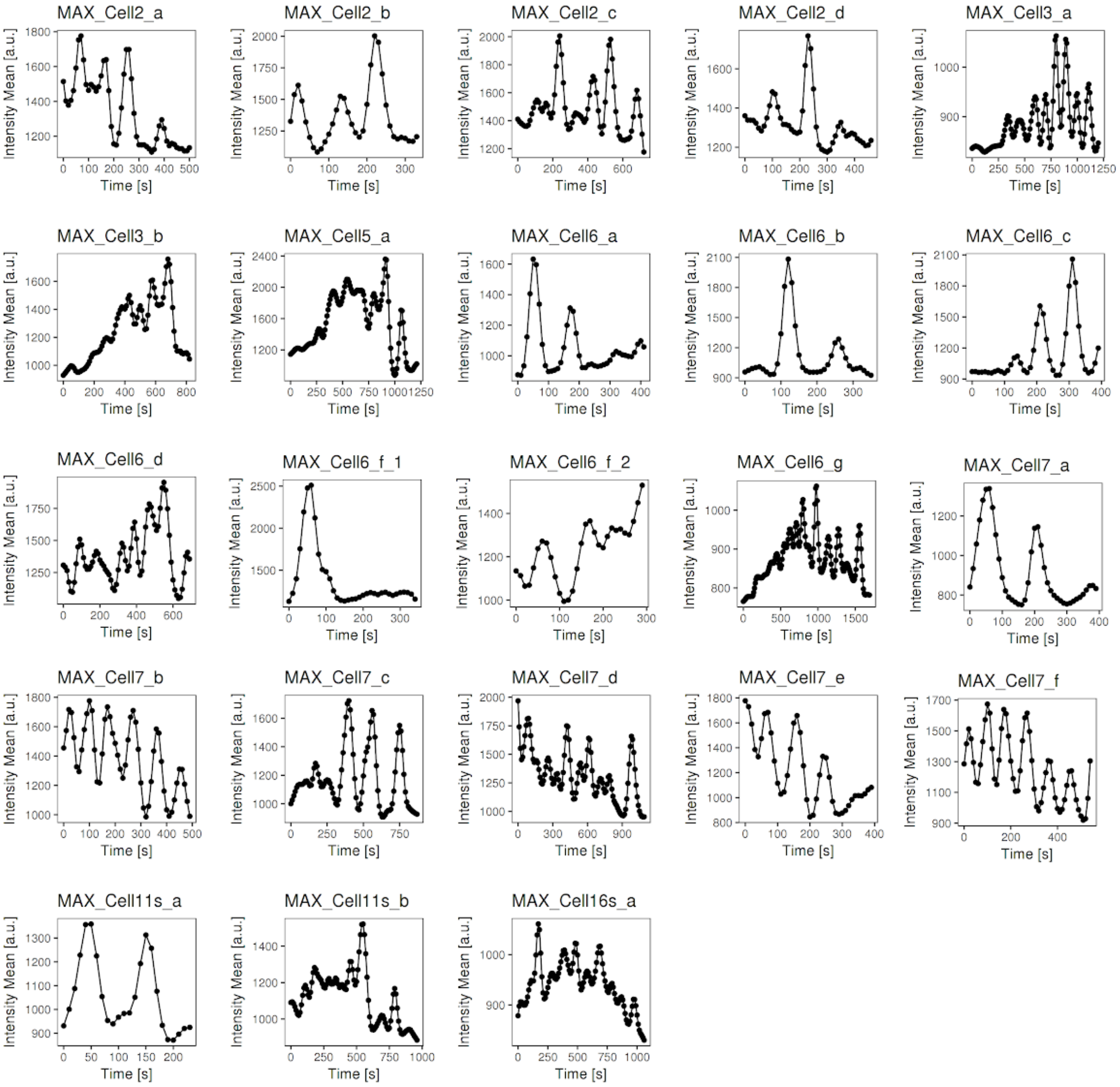

**Supplementary Figure S6. Splined quantified time courses for ATG13 for the mitophagy model.**

To facilitate the synchronisation, ATG13 time courses were splined in order to remove small noise in the signal and visualise the peaks more clearly. The splining of this data was performed using the R function smooth.spline() with spline parameter *spar* set to 0.4.

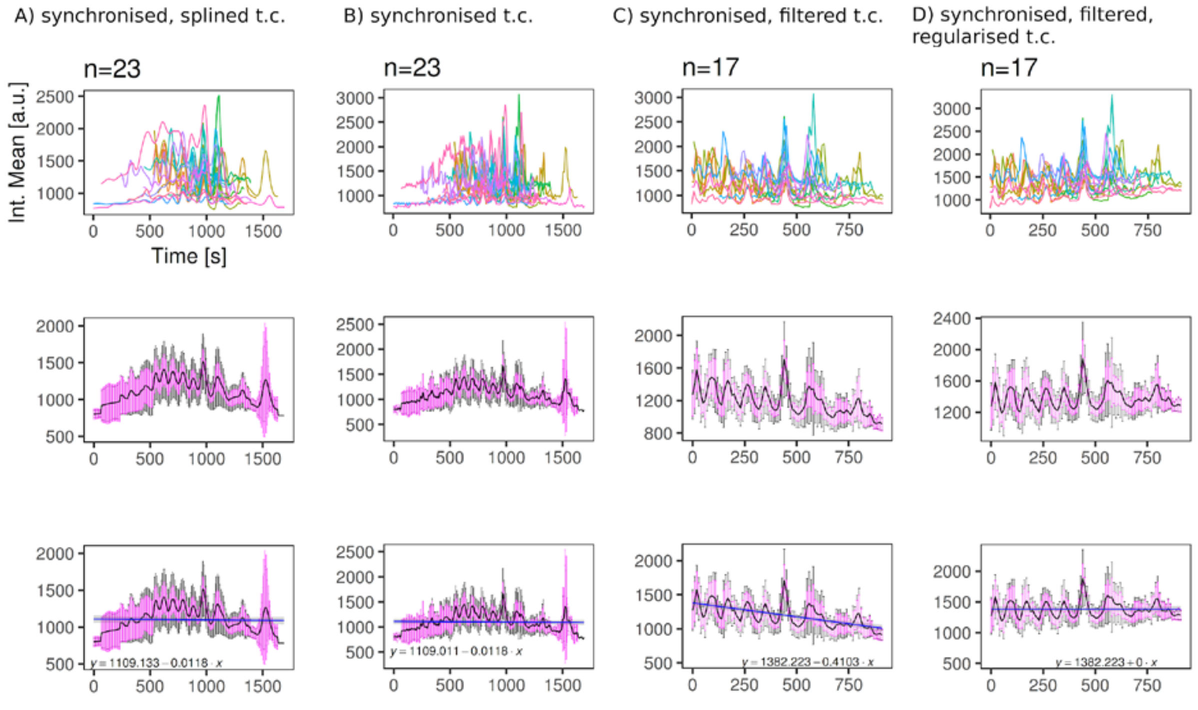

**Supplementary Figure S7. Synchronisation and filtering of ATG13 time courses for the mitophagy model.**

A) ATG13 splined time courses were synchronised by overlapping the delays between peaks.
B) The splined time courses were then replaced with the original ATG13 time courses.
C) Six time courses were removed because they contained too much noise in their signal. Initial and later signals were cut off in order to have at least 3 repeats for each time point.
D) The mean time course was regularised so that there was no signal decline over time. Each time course was adjusted based on this regularisation.

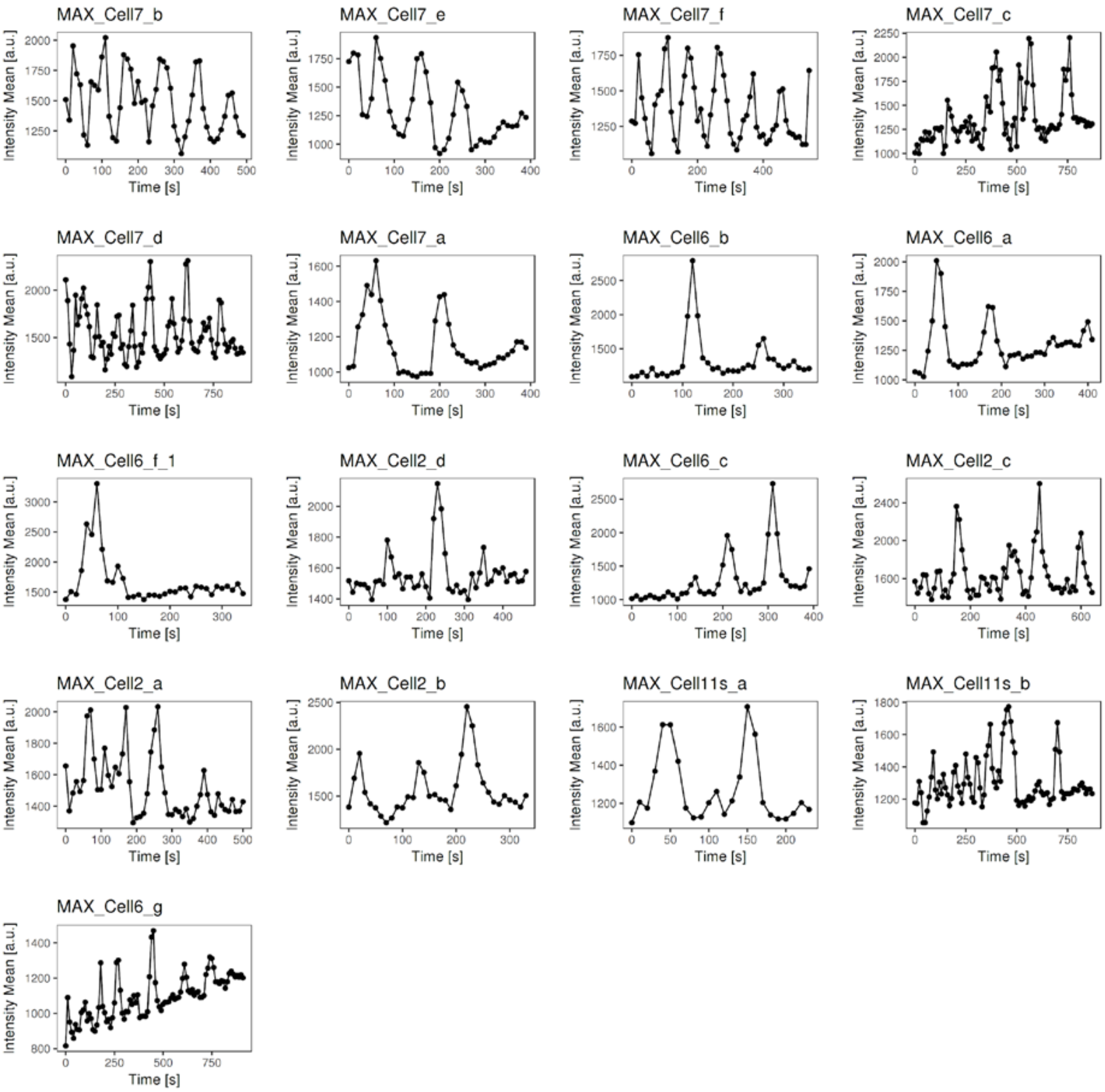

**Supplementary Figure S8. ATG13 time courses for the mitophagy model after synchronisation, filtering and regularisation.**

ATG13 final time courses are illustrated. The data points from this figure were used for parameter estimation of the mitophagy model.

**Supplementary Model 1. Non-selective autophagy model 3.**

**Supplementary Model 2. Mitophagy model.**

```
<?xml version="1.0" encoding="UTF-8"?>
<!-- Created by COPASI version 4.22 (Build 170) on 2018-07-12 13:21 with
libSBML version 5.15.3. -->
<sbml xmlns="http://www.sbml.org/sbml/level3/version1/core"
xmlns:layout="http://www.sbml.org/sbml/level3/version1/layout/version1
"
xmlns:render="http://www.sbml.org/sbml/level3/version1/render/version1
" level="3" version="1" layout:required="false" render:required="false">
  <model metaid="COPASI0" id="m19" name="m19" substanceUnits="substance"
timeUnits="time" volumeUnits="volume" areaUnits="area"
lengthUnits="length" extentUnits="substance">
    <annotation>
      <COPASI xmlns="http://www.copasi.org/static/sbml">
        <rdf:RDF xmlns:dcterms="http://purl.org/dc/terms/"
xmlns:rdf="http://www.w3.org/1999/02/22-rdf-syntax-ns#">
          <rdf:Description rdf:about="#COPASI0">
            <dcterms:created>
              <rdf:Description>
                <dcterms:W3CDTF>2013-07-23T11:01:00Z</dcterms:W3CDTF>
              </rdf:Description>
            </dcterms:created>
          </rdf:Description>
        </rdf:RDF>
      </COPASI>
    </annotation>
    <listOfFunctionDefinitions>
      <functionDefinition metaid="COPASI22" id="f_prod" name="f_prod">
        <annotation>
          <COPASI xmlns="http://www.copasi.org/static/sbml">
            <rdf:RDF xmlns:dcterms="http://purl.org/dc/terms/"
xmlns:rdf="http://www.w3.org/1999/02/22-rdf-syntax-ns#">
              <rdf:Description rdf:about="#COPASI22">
                <dcterms:created>
                  <rdf:Description>

<dcterms:W3CDTF>2017-11-22T13:35:42Z</dcterms:W3CDTF>
                  </rdf:Description>
                </dcterms:created>
              </rdf:Description>
            </rdf:RDF>
          </COPASI>
        </annotation>
        <math xmlns="http://www.w3.org/1998/Math/MathML">
          <lambda>
            <bvar>
              <ci> k </ci>
            </bvar>
            <bvar>
              <ci> A </ci>
            </bvar>
            <bvar>
              <ci> m </ci>
```

```
            &lt;/bvar&gt;
            &lt;bvar&gt;
              &lt;ci&gt; w &lt;/ci&gt;
            &lt;/bvar&gt;
            &lt;bvar&gt;
              &lt;ci&gt; wk &lt;/ci&gt;
            &lt;/bvar&gt;
            &lt;bvar&gt;
              &lt;ci&gt; n &lt;/ci&gt;
            &lt;/bvar&gt;
            &lt;apply&gt;
              &lt;divide/&gt;
              &lt;apply&gt;
                &lt;times/&gt;
                &lt;ci&gt; k &lt;/ci&gt;
                &lt;apply&gt;
                  &lt;power/&gt;
                  &lt;ci&gt; A &lt;/ci&gt;
                  &lt;ci&gt; m &lt;/ci&gt;
                &lt;/apply&gt;
              &lt;/apply&gt;
              &lt;apply&gt;
                &lt;plus/&gt;
                &lt;cn&gt; 1 &lt;/cn&gt;
                &lt;apply&gt;
                  &lt;power/&gt;
                  &lt;apply&gt;
                    &lt;divide/&gt;
                    &lt;ci&gt; w &lt;/ci&gt;
                    &lt;ci&gt; wk &lt;/ci&gt;
                  &lt;/apply&gt;
                  &lt;ci&gt; n &lt;/ci&gt;
                &lt;/apply&gt;
              &lt;/apply&gt;
            &lt;/apply&gt;
          &lt;/lambda&gt;
        &lt;/math&gt;
      &lt;/functionDefinition&gt;
      &lt;functionDefinition metaid="COPASI23" id="f_rem" name="f_rem"&gt;
        &lt;annotation&gt;
          &lt;COPASI xmlns="http://www.copasi.org/static/sbml"&gt;
            <rdf:RDF xmlns:dcterms="http://purl.org/dc/terms/"
xmlns:rdf="http://www.w3.org/1999/02/22-rdf-syntax-ns#">
              &lt;rdf:Description rdf:about="#COPASI23"&gt;
                &lt;dcterms:created&gt;
                  &lt;rdf:Description&gt;

&lt;dcterms:W3CDTF&gt;2017-11-22T13:35:42Z&lt;/dcterms:W3CDTF&gt;
                  &lt;/rdf:Description&gt;
                &lt;/dcterms:created&gt;
              &lt;/rdf:Description&gt;
            &lt;/rdf:RDF&gt;
```

```
        </COPASI>
      </annotation>
      <math xmlns="http://www.w3.org/1998/Math/MathML">
        <lambda>
          <bvar>
            <ci> k </ci>
          </bvar>
          <bvar>
            <ci> A </ci>
          </bvar>
          <bvar>
            <ci> m </ci>
          </bvar>
          <bvar>
            <ci> w </ci>
          </bvar>
          <bvar>
            <ci> wk </ci>
          </bvar>
          <bvar>
            <ci> n </ci>
          </bvar>
          <apply>
            <times/>
            <ci> k </ci>
            <apply>
              <power/>
              <ci> A </ci>
              <apply>
                <plus/>
                <cn> 1 </cn>
                <ci> m </ci>
              </apply>
            </apply>
            <apply>
              <plus/>
              <cn> 1 </cn>
              <apply>
                <power/>
                <apply>
                  <divide/>
                  <ci> w </ci>
                  <ci> wk </ci>
                </apply>
                <ci> n </ci>
              </apply>
            </apply>
          </apply>
        </lambda>
      </math>
    </functionDefinition>
  </listOfFunctionDefinitions>
```

```
    <listOfUnitDefinitions>
      <unitDefinition id="length" name="length">
        <listOfUnits>
          <unit kind="metre" exponent="1" scale="0" multiplier="1"/>
        </listOfUnits>
      </unitDefinition>
      <unitDefinition id="area" name="area">
        <listOfUnits>
          <unit kind="metre" exponent="2" scale="0" multiplier="1"/>
        </listOfUnits>
      </unitDefinition>
      <unitDefinition id="volume" name="volume">
        <listOfUnits>
          <unit kind="litre" exponent="1" scale="-3" multiplier="1"/>
        </listOfUnits>
      </unitDefinition>
      <unitDefinition id="time" name="time">
        <listOfUnits>
          <unit kind="second" exponent="1" scale="0" multiplier="1"/>
        </listOfUnits>
      </unitDefinition>
      <unitDefinition id="substance" name="substance">
        <listOfUnits>
          <unit kind="mole" exponent="1" scale="-3" multiplier="1"/>
        </listOfUnits>
      </unitDefinition>
    </listOfUnitDefinitions>
    <listOfCompartments>
      <compartment metaid="COPASI1" id="ER" name="ER"
spatialDimensions="3" size="1" units="volume" constant="true">
        <annotation>
          <COPASI xmlns="http://www.copasi.org/static/sbml">
            <rdf:RDF xmlns:dcterms="http://purl.org/dc/terms/"
xmlns:rdf="http://www.w3.org/1999/02/22-rdf-syntax-ns#">
              <rdf:Description rdf:about="#COPASI1">
                <dcterms:created>
                  <rdf:Description>

<dcterms:W3CDTF>2017-11-22T13:32:41Z</dcterms:W3CDTF>
                  </rdf:Description>
                </dcterms:created>
              </rdf:Description>
            </rdf:RDF>
          </COPASI>
        </annotation>
      </compartment>
    </listOfCompartments>
    <listOfSpecies>
      <species metaid="COPASI2" id="ATG13" name="ATG13" compartment="ER"
initialConcentration="1" substanceUnits="substance"
hasOnlySubstanceUnits="false" boundaryCondition="false"
constant="false">
```

```
        <annotation>
          <COPASI xmlns="http://www.copasi.org/static/sbml">
            <rdf:RDF xmlns:dcterms="http://purl.org/dc/terms/"
xmlns:rdf="http://www.w3.org/1999/02/22-rdf-syntax-ns#">
              <rdf:Description rdf:about="#COPASI2">
                <dcterms:created>
                  <rdf:Description>

<dcterms:W3CDTF>2017-11-22T13:32:48Z</dcterms:W3CDTF>
                  </rdf:Description>
                </dcterms:created>
              </rdf:Description>
            </rdf:RDF>
          </COPASI>
        </annotation>
      </species>
    </listOfSpecies>
    <listOfParameters>
      <parameter metaid="COPASI3" id="kprodATG13" name="kprodATG13"
value="0.00822279" constant="false">
        <annotation>
          <COPASI xmlns="http://www.copasi.org/static/sbml">
            <rdf:RDF xmlns:dcterms="http://purl.org/dc/terms/"
xmlns:rdf="http://www.w3.org/1999/02/22-rdf-syntax-ns#">
              <rdf:Description rdf:about="#COPASI3">
                <dcterms:created>
                  <rdf:Description>

<dcterms:W3CDTF>2016-09-20T16:37:39Z</dcterms:W3CDTF>
                  </rdf:Description>
                </dcterms:created>
              </rdf:Description>
            </rdf:RDF>
          </COPASI>
        </annotation>
      </parameter>
      <parameter metaid="COPASI4" id="ATG13_obs" name="ATG13_obs"
value="525" constant="false">
        <annotation>
          <COPASI xmlns="http://www.copasi.org/static/sbml">
            <rdf:RDF xmlns:dcterms="http://purl.org/dc/terms/"
xmlns:rdf="http://www.w3.org/1999/02/22-rdf-syntax-ns#">
              <rdf:Description rdf:about="#COPASI4">
                <dcterms:created>
                  <rdf:Description>

<dcterms:W3CDTF>2016-09-22T10:52:54Z</dcterms:W3CDTF>
                  </rdf:Description>
                </dcterms:created>
              </rdf:Description>
            </rdf:RDF>
          </COPASI>
```

```
        </annotation>
      </parameter>
      <parameter metaid="COPASI5" id="wrtm" name="wrtm" value="0"
constant="false">
        <annotation>
          <COPASI xmlns="http://www.copasi.org/static/sbml">
            <rdf:RDF xmlns:dcterms="http://purl.org/dc/terms/"
xmlns:rdf="http://www.w3.org/1999/02/22-rdf-syntax-ns#">
              <rdf:Description rdf:about="#COPASI5">
                <dcterms:created>
                  <rdf:Description>

<dcterms:W3CDTF>2016-09-22T15:46:47Z</dcterms:W3CDTF>
                  </rdf:Description>
                </dcterms:created>
              </rdf:Description>
            </rdf:RDF>
          </COPASI>
        </annotation>
      </parameter>
      <parameter metaid="COPASI6" id="wrtm_flag" name="wrtm_flag"
value="0" constant="true">
        <annotation>
          <COPASI xmlns="http://www.copasi.org/static/sbml">
            <rdf:RDF xmlns:dcterms="http://purl.org/dc/terms/"
xmlns:rdf="http://www.w3.org/1999/02/22-rdf-syntax-ns#">
              <rdf:Description rdf:about="#COPASI6">
                <dcterms:created>
                  <rdf:Description>

<dcterms:W3CDTF>2016-09-22T15:46:39Z</dcterms:W3CDTF>
                  </rdf:Description>
                </dcterms:created>
              </rdf:Description>
            </rdf:RDF>
          </COPASI>
        </annotation>
      </parameter>
      <parameter metaid="COPASI7" id="kwrtm" name="kwrtm"
value="1.73235" constant="true">
        <annotation>
          <COPASI xmlns="http://www.copasi.org/static/sbml">
            <rdf:RDF xmlns:dcterms="http://purl.org/dc/terms/"
xmlns:rdf="http://www.w3.org/1999/02/22-rdf-syntax-ns#">
              <rdf:Description rdf:about="#COPASI7">
                <dcterms:created>
                  <rdf:Description>

<dcterms:W3CDTF>2016-09-22T15:48:18Z</dcterms:W3CDTF>
                  </rdf:Description>
                </dcterms:created>
              </rdf:Description>
```

```
            </rdf:RDF>
          </COPASI>
        </annotation>
      </parameter>
      <parameter metaid="COPASI8" id="ATG13_sf" name="ATG13_sf"
value="525" constant="true">
        <annotation>
          <COPASI xmlns="http://www.copasi.org/static/sbml">
            <rdf:RDF xmlns:dcterms="http://purl.org/dc/terms/"
xmlns:rdf="http://www.w3.org/1999/02/22-rdf-syntax-ns#">
              <rdf:Description rdf:about="#COPASI8">
                <dcterms:created>
                  <rdf:Description>

<dcterms:W3CDTF>2016-09-27T15:55:01Z</dcterms:W3CDTF>
                  </rdf:Description>
                </dcterms:created>
              </rdf:Description>
            </rdf:RDF>
          </COPASI>
        </annotation>
      </parameter>
      <parameter metaid="COPASI9" id="ATG13_ini" name="ATG13_ini"
value="525" constant="true">
        <annotation>
          <COPASI xmlns="http://www.copasi.org/static/sbml">
            <rdf:RDF xmlns:dcterms="http://purl.org/dc/terms/"
xmlns:rdf="http://www.w3.org/1999/02/22-rdf-syntax-ns#">
              <rdf:Description rdf:about="#COPASI9">
                <dcterms:created>
                  <rdf:Description>

<dcterms:W3CDTF>2016-09-27T16:49:27Z</dcterms:W3CDTF>
                  </rdf:Description>
                </dcterms:created>
              </rdf:Description>
            </rdf:RDF>
          </COPASI>
        </annotation>
      </parameter>
      <parameter metaid="COPASI10" id="kremATG13" name="kremATG13"
value="0.00284586" constant="false">
        <annotation>
          <COPASI xmlns="http://www.copasi.org/static/sbml">
            <rdf:RDF xmlns:dcterms="http://purl.org/dc/terms/"
xmlns:rdf="http://www.w3.org/1999/02/22-rdf-syntax-ns#">
              <rdf:Description rdf:about="#COPASI10">
                <dcterms:created>
                  <rdf:Description>

<dcterms:W3CDTF>2016-10-17T11:48:37Z</dcterms:W3CDTF>
                  </rdf:Description>
```

```
                </dcterms:created>
              </rdf:Description>
            </rdf:RDF>
          </COPASI>
        </annotation>
      </parameter>
      <parameter metaid="COPASI11" id="n" name="n" value="1"
constant="true">
        <annotation>
          <COPASI xmlns="http://www.copasi.org/static/sbml">
            <rdf:RDF xmlns:dcterms="http://purl.org/dc/terms/"
xmlns:rdf="http://www.w3.org/1999/02/22-rdf-syntax-ns#">
              <rdf:Description rdf:about="#COPASI11">
                <dcterms:created>
                  <rdf:Description>

<dcterms:W3CDTF>2016-10-21T14:26:01Z</dcterms:W3CDTF>
                  </rdf:Description>
                </dcterms:created>
              </rdf:Description>
            </rdf:RDF>
          </COPASI>
        </annotation>
      </parameter>
      <parameter metaid="COPASI12" id="t" name="t" value="225.454"
constant="true">
        <annotation>
          <COPASI xmlns="http://www.copasi.org/static/sbml">
            <rdf:RDF xmlns:dcterms="http://purl.org/dc/terms/"
xmlns:rdf="http://www.w3.org/1999/02/22-rdf-syntax-ns#">
              <rdf:Description rdf:about="#COPASI12">
                <dcterms:created>
                  <rdf:Description>

<dcterms:W3CDTF>2016-10-21T16:51:48Z</dcterms:W3CDTF>
                  </rdf:Description>
                </dcterms:created>
              </rdf:Description>
            </rdf:RDF>
          </COPASI>
        </annotation>
      </parameter>
      <parameter metaid="COPASI13" id="wrtm_si" name="wrtm_si"
value="1500" constant="true">
        <annotation>
          <COPASI xmlns="http://www.copasi.org/static/sbml">
            <rdf:RDF xmlns:dcterms="http://purl.org/dc/terms/"
xmlns:rdf="http://www.w3.org/1999/02/22-rdf-syntax-ns#">
              <rdf:Description rdf:about="#COPASI13">
                <dcterms:created>
                  <rdf:Description>
```

```
<dcterms:W3CDTF>2016-10-26T16:07:58Z</dcterms:W3CDTF>
                  </rdf:Description>
                </dcterms:created>
              </rdf:Description>
            </rdf:RDF>
          </COPASI>
        </annotation>
      </parameter>
      <parameter metaid="COPASI14" id="wrtm_sf" name="wrtm_sf"
value="1500" constant="true">
        <annotation>
          <COPASI xmlns="http://www.copasi.org/static/sbml">
            <rdf:RDF xmlns:dcterms="http://purl.org/dc/terms/"
xmlns:rdf="http://www.w3.org/1999/02/22-rdf-syntax-ns#">
              <rdf:Description rdf:about="#COPASI14">
                <dcterms:created>
                  <rdf:Description>

<dcterms:W3CDTF>2016-10-26T16:08:06Z</dcterms:W3CDTF>
                  </rdf:Description>
                </dcterms:created>
              </rdf:Description>
            </rdf:RDF>
          </COPASI>
        </annotation>
      </parameter>
      <parameter metaid="COPASI15" id="wrtm_scaled" name="wrtm_scaled"
value="0" constant="false">
        <annotation>
          <COPASI xmlns="http://www.copasi.org/static/sbml">
            <rdf:RDF xmlns:dcterms="http://purl.org/dc/terms/"
xmlns:rdf="http://www.w3.org/1999/02/22-rdf-syntax-ns#">
              <rdf:Description rdf:about="#COPASI15">
                <dcterms:created>
                  <rdf:Description>

<dcterms:W3CDTF>2016-10-26T16:08:18Z</dcterms:W3CDTF>
                  </rdf:Description>
                </dcterms:created>
              </rdf:Description>
            </rdf:RDF>
          </COPASI>
        </annotation>
      </parameter>
      <parameter metaid="COPASI16" id="m" name="m" value="1.01365"
constant="true">
        <annotation>
          <COPASI xmlns="http://www.copasi.org/static/sbml">
            <rdf:RDF xmlns:dcterms="http://purl.org/dc/terms/"
xmlns:rdf="http://www.w3.org/1999/02/22-rdf-syntax-ns#">
              <rdf:Description rdf:about="#COPASI16">
                <dcterms:created>
```

```
                  <rdf:Description>
<dcterms:W3CDTF>2016-10-27T12:48:31Z</dcterms:W3CDTF>
                  </rdf:Description>
                </dcterms:created>
              </rdf:Description>
            </rdf:RDF>
          </COPASI>
        </annotation>
      </parameter>
      <parameter id="ModelValue_0" name="Initial for kprodATG13"
value="0.00822279" constant="true">
        <annotation>
          <initialValue xmlns="http://copasi.org/initialValue"
parent="kprodATG13"/>
        </annotation>
      </parameter>
      <parameter id="ModelValue_7" name="Initial for kremATG13"
value="0.00284586" constant="true">
        <annotation>
          <initialValue xmlns="http://copasi.org/initialValue"
parent="kremATG13"/>
        </annotation>
      </parameter>
    </listOfParameters>
    <listOfInitialAssignments>
      <initialAssignment symbol="ATG13">
        <math xmlns="http://www.w3.org/1998/Math/MathML">
          <apply>
            <divide/>
            <ci> ATG13_ini </ci>
            <ci> ATG13_sf </ci>
          </apply>
        </math>
      </initialAssignment>
      <initialAssignment symbol="ModelValue_0">
        <math xmlns="http://www.w3.org/1998/Math/MathML">
          <ci> kprodATG13 </ci>
        </math>
      </initialAssignment>
      <initialAssignment symbol="ModelValue_7">
        <math xmlns="http://www.w3.org/1998/Math/MathML">
          <ci> kremATG13 </ci>
        </math>
      </initialAssignment>
    </listOfInitialAssignments>
    <listOfRules>
      <assignmentRule variable="ATG13_obs">
        <math xmlns="http://www.w3.org/1998/Math/MathML">
          <apply>
            <times/>
            <ci> ATG13_sf </ci>
```

```
            <ci> ATG13 </ci>
          </apply>
        </math>
      </assignmentRule>
      <assignmentRule variable="wrtm">
        <math xmlns="http://www.w3.org/1998/Math/MathML">
          <piecewise>
            <piece>
              <apply>
                <divide/>
                <cn> 0 </cn>
                <ci> wrtm_sf </ci>
              </apply>
              <apply>
                <eq/>
                <ci> wrtm_flag </ci>
                <cn> 0 </cn>
              </apply>
            </piece>
            <otherwise>
              <apply>
                <divide/>
                <ci> wrtm_si </ci>
                <ci> wrtm_sf </ci>
              </apply>
            </otherwise>
          </piecewise>
        </math>
      </assignmentRule>
      <assignmentRule variable="wrtm_scaled">
        <math xmlns="http://www.w3.org/1998/Math/MathML">
          <apply>
            <times/>
            <ci> wrtm_sf </ci>
            <ci> wrtm </ci>
          </apply>
        </math>
      </assignmentRule>
    </listOfRules>
    <listOfReactions>
      <reaction metaid="COPASI20" id="ATG13_prod" name="ATG13_prod"
reversible="false" fast="false">
        <annotation>
          <COPASI xmlns="http://www.copasi.org/static/sbml">
            <rdf:RDF xmlns:dcterms="http://purl.org/dc/terms/"
xmlns:rdf="http://www.w3.org/1999/02/22-rdf-syntax-ns#">
              <rdf:Description rdf:about="#COPASI20">
                <dcterms:created>
                  <rdf:Description>

<dcterms:W3CDTF>2017-11-22T13:38:36Z</dcterms:W3CDTF>
                  </rdf:Description>
```

```
              &lt;/dcterms:created&gt;
            &lt;/rdf:Description&gt;
          &lt;/rdf:RDF&gt;
        &lt;/COPASI&gt;
      &lt;/annotation&gt;
      &lt;listOfProducts&gt;
        <speciesReference species="ATG13" stoichiometry="1"
constant="true"/>
      &lt;/listOfProducts&gt;
      &lt;kineticLaw&gt;
        &lt;math xmlns="http://www.w3.org/1998/Math/MathML"&gt;
          &lt;apply&gt;
            &lt;times/&gt;
            &lt;ci&gt; ER &lt;/ci&gt;
            &lt;apply&gt;
              &lt;ci&gt; f_prod &lt;/ci&gt;
              &lt;ci&gt; kprodATG13 &lt;/ci&gt;
              &lt;ci&gt; ATG13 &lt;/ci&gt;
              &lt;ci&gt; m &lt;/ci&gt;
              &lt;ci&gt; wrtm &lt;/ci&gt;
              &lt;ci&gt; kwrtm &lt;/ci&gt;
              &lt;ci&gt; n &lt;/ci&gt;
            &lt;/apply&gt;
          &lt;/apply&gt;
        &lt;/math&gt;
      &lt;/kineticLaw&gt;
    &lt;/reaction&gt;
    <reaction metaid="COPASI21" id="ATG13_rem" name="ATG13_rem"
reversible="false" fast="false">
      &lt;annotation&gt;
        &lt;COPASI xmlns="http://www.copasi.org/static/sbml"&gt;
          <rdf:RDF xmlns:dcterms="http://purl.org/dc/terms/"
xmlns:rdf="http://www.w3.org/1999/02/22-rdf-syntax-ns#">
            &lt;rdf:Description rdf:about="#COPASI21"&gt;
              &lt;dcterms:created&gt;
                &lt;rdf:Description&gt;

&lt;dcterms:W3CDTF&gt;2017-11-22T13:43:39Z&lt;/dcterms:W3CDTF&gt;
                &lt;/rdf:Description&gt;
              &lt;/dcterms:created&gt;
            &lt;/rdf:Description&gt;
          &lt;/rdf:RDF&gt;
        &lt;/COPASI&gt;
      &lt;/annotation&gt;
      &lt;listOfReactants&gt;
        <speciesReference species="ATG13" stoichiometry="1"
constant="true"/>
      &lt;/listOfReactants&gt;
      &lt;kineticLaw&gt;
        &lt;math xmlns="http://www.w3.org/1998/Math/MathML"&gt;
          &lt;apply&gt;
            &lt;times/&gt;
```

```
              <ci> ER </ci>
              <apply>
                <ci> f_rem </ci>
                <ci> kremATG13 </ci>
                <ci> ATG13 </ci>
                <ci> m </ci>
                <ci> wrtm </ci>
                <ci> kwrtm </ci>
                <ci> n </ci>
              </apply>
            </apply>
          </math>
        </kineticLaw>
      </reaction>
    </listOfReactions>
    <listOfEvents>
      <event metaid="COPASI17" id="atg13_accumulation"
name="atg13_accumulation" useValuesFromTriggerTime="false">
        <annotation>
          <COPASI xmlns="http://www.copasi.org/static/sbml">
            <rdf:RDF xmlns:dcterms="http://purl.org/dc/terms/"
xmlns:rdf="http://www.w3.org/1999/02/22-rdf-syntax-ns#">
              <rdf:Description rdf:about="#COPASI17">
                <dcterms:created>
                  <rdf:Description>

<dcterms:W3CDTF>2017-11-22T13:22:45Z</dcterms:W3CDTF>
                  </rdf:Description>
                </dcterms:created>
              </rdf:Description>
            </rdf:RDF>
          </COPASI>
        </annotation>
        <trigger initialValue="false" persistent="true">
          <math xmlns="http://www.w3.org/1998/Math/MathML">
            <apply>
              <lt/>
              <csymbol encoding="text"
definitionURL="http://www.sbml.org/sbml/symbols/time"> time </csymbol>
              <ci> t </ci>
            </apply>
          </math>
        </trigger>
        <listOfEventAssignments>
          <eventAssignment variable="kprodATG13">
            <math xmlns="http://www.w3.org/1998/Math/MathML">
              <ci> ModelValue_0 </ci>
            </math>
          </eventAssignment>
          <eventAssignment variable="kremATG13">
            <math xmlns="http://www.w3.org/1998/Math/MathML">
              <cn> 0 </cn>
```

```
            </math>
          </eventAssignment>
        </listOfEventAssignments>
      </event>
      <event metaid="COPASI18" id="atg13_removal" name="atg13_removal"
useValuesFromTriggerTime="false">
        <annotation>
          <COPASI xmlns="http://www.copasi.org/static/sbml">
            <rdf:RDF xmlns:dcterms="http://purl.org/dc/terms/"
xmlns:rdf="http://www.w3.org/1999/02/22-rdf-syntax-ns#">
              <rdf:Description rdf:about="#COPASI18">
                <dcterms:created>
                  <rdf:Description>

<dcterms:W3CDTF>2017-11-22T13:22:45Z</dcterms:W3CDTF>
                  </rdf:Description>
                </dcterms:created>
              </rdf:Description>
            </rdf:RDF>
          </COPASI>
        </annotation>
        <trigger initialValue="true" persistent="true">
          <math xmlns="http://www.w3.org/1998/Math/MathML">
            <apply>
              <and/>
              <apply>
                <gt/>
                <ci> kprodATG13 </ci>
                <cn> 0 </cn>
              </apply>
              <apply>
                <eq/>
                <ci> kremATG13 </ci>
                <cn> 0 </cn>
              </apply>
            </apply>
          </math>
        </trigger>
        <delay>
          <math xmlns="http://www.w3.org/1998/Math/MathML">
            <ci> t </ci>
          </math>
        </delay>
        <listOfEventAssignments>
          <eventAssignment variable="kprodATG13">
            <math xmlns="http://www.w3.org/1998/Math/MathML">
              <cn> 0 </cn>
            </math>
          </eventAssignment>
          <eventAssignment variable="kremATG13">
            <math xmlns="http://www.w3.org/1998/Math/MathML">
              <ci> ModelValue_7 </ci>
```

```
            </math>
          </eventAssignment>
        </listOfEventAssignments>
      </event>
      <event metaid="COPASI19" id="atg13_basal" name="atg13_basal"
useValuesFromTriggerTime="false">
        <annotation>
          <COPASI xmlns="http://www.copasi.org/static/sbml">
            <rdf:RDF xmlns:dcterms="http://purl.org/dc/terms/"
xmlns:rdf="http://www.w3.org/1999/02/22-rdf-syntax-ns#">
              <rdf:Description rdf:about="#COPASI19">
                <dcterms:created>
                  <rdf:Description>

<dcterms:W3CDTF>2017-11-22T13:50:30Z</dcterms:W3CDTF>
                  </rdf:Description>
                </dcterms:created>
              </rdf:Description>
            </rdf:RDF>
          </COPASI>
        </annotation>
        <trigger initialValue="true" persistent="true">
          <math xmlns="http://www.w3.org/1998/Math/MathML">
            <apply>
              <and/>
              <apply>
                <and/>
                <apply>
                  <eq/>
                  <ci> kprodATG13 </ci>
                  <cn> 0 </cn>
                </apply>
                <apply>
                  <gt/>
                  <ci> kremATG13 </ci>
                  <cn> 0 </cn>
                </apply>
              </apply>
              <apply>
                <lt/>
                <ci> ATG13 </ci>
                <cn> 1 </cn>
              </apply>
            </apply>
          </math>
        </trigger>
        <listOfEventAssignments>
          <eventAssignment variable="kprodATG13">
            <math xmlns="http://www.w3.org/1998/Math/MathML">
              <cn> 0 </cn>
            </math>
          </eventAssignment>
```

```
          <eventAssignment variable="kremATG13">
            <math xmlns="http://www.w3.org/1998/Math/MathML">
              <cn> 0 </cn>
            </math>
          </eventAssignment>
        </listOfEventAssignments>
      </event>
    </listOfEvents>
  </model>
</sbml>
```

```
<?xml version="1.0" encoding="UTF-8"?>
<!-- Created by COPASI version 4.22 (Build 170) on 2018-07-12 13:21 with
libSBML version 5.15.3. -->
<sbml xmlns="http://www.sbml.org/sbml/level3/version1/core"
xmlns:layout="http://www.sbml.org/sbml/level3/version1/layout/version1
"
xmlns:render="http://www.sbml.org/sbml/level3/version1/render/version1
" level="3" version="1" layout:required="false" render:required="false">
  <model metaid="COPASI0" id="mitophagy_ATG13_model" name="mitophagy
ATG13 model" substanceUnits="substance" timeUnits="time"
volumeUnits="volume" areaUnits="area" lengthUnits="length"
extentUnits="substance">
    <annotation>
      <COPASI xmlns="http://www.copasi.org/static/sbml">
        <rdf:RDF xmlns:dcterms="http://purl.org/dc/terms/"
xmlns:rdf="http://www.w3.org/1999/02/22-rdf-syntax-ns#">
          <rdf:Description rdf:about="#COPASI0">
            <dcterms:created>
              <rdf:Description>
                <dcterms:W3CDTF>2013-07-23T11:01:00Z</dcterms:W3CDTF>
              </rdf:Description>
            </dcterms:created>
          </rdf:Description>
        </rdf:RDF>
      </COPASI>
    </annotation>
    <listOfFunctionDefinitions>
      <functionDefinition id="RNORMAL">
        <annotation>
          <distribution
xmlns="http://sbml.org/annotations/distribution"
definition="http://www.uncertml.org/distributions/normal"/>
        </annotation>
        <math xmlns="http://www.w3.org/1998/Math/MathML">
          <lambda>
            <bvar>
              <ci> m </ci>
            </bvar>
            <bvar>
              <ci> s </ci>
            </bvar>
            <ci> m </ci>
          </lambda>
        </math>
      </functionDefinition>
      <functionDefinition metaid="COPASI36" id="f_prod_atg13"
name="f_prod_atg13">
        <annotation>
          <COPASI xmlns="http://www.copasi.org/static/sbml">
            <rdf:RDF xmlns:dcterms="http://purl.org/dc/terms/"
xmlns:rdf="http://www.w3.org/1999/02/22-rdf-syntax-ns#">
              <rdf:Description rdf:about="#COPASI36">
```

```
                  <dcterms:created>
                    <rdf:Description>

<dcterms:W3CDTF>2017-11-22T13:35:42Z</dcterms:W3CDTF>
                    </rdf:Description>
                  </dcterms:created>
                </rdf:Description>
              </rdf:RDF>
            </COPASI>
          </annotation>
          <math xmlns="http://www.w3.org/1998/Math/MathML">
            <lambda>
              <bvar>
                <ci> k </ci>
              </bvar>
              <bvar>
                <ci> A </ci>
              </bvar>
              <bvar>
                <ci> m </ci>
              </bvar>
              <bvar>
                <ci> w </ci>
              </bvar>
              <bvar>
                <ci> wk </ci>
              </bvar>
              <bvar>
                <ci> n </ci>
              </bvar>
              <apply>
                <divide/>
                <apply>
                  <times/>
                  <ci> k </ci>
                  <apply>
                    <power/>
                    <ci> A </ci>
                    <ci> m </ci>
                  </apply>
                </apply>
                <apply>
                  <plus/>
                  <cn> 1 </cn>
                  <apply>
                    <power/>
                    <apply>
                      <divide/>
                      <ci> w </ci>
                      <ci> wk </ci>
                    </apply>
                    <ci> n </ci>
```

```
              </apply>
            </apply>
          </apply>
        </lambda>
      </math>
    </functionDefinition>
    <functionDefinition metaid="COPASI37" id="f_rem_atg13"
name="f_rem_atg13">
      <annotation>
        <COPASI xmlns="http://www.copasi.org/static/sbml">
          <rdf:RDF xmlns:dcterms="http://purl.org/dc/terms/"
xmlns:rdf="http://www.w3.org/1999/02/22-rdf-syntax-ns#">
            <rdf:Description rdf:about="#COPASI37">
              <dcterms:created>
                <rdf:Description>

<dcterms:W3CDTF>2017-11-22T13:35:42Z</dcterms:W3CDTF>
                </rdf:Description>
              </dcterms:created>
            </rdf:Description>
          </rdf:RDF>
        </COPASI>
      </annotation>
      <math xmlns="http://www.w3.org/1998/Math/MathML">
        <lambda>
          <bvar>
            <ci> k </ci>
          </bvar>
          <bvar>
            <ci> A </ci>
          </bvar>
          <bvar>
            <ci> m </ci>
          </bvar>
          <bvar>
            <ci> w </ci>
          </bvar>
          <bvar>
            <ci> wk </ci>
          </bvar>
          <bvar>
            <ci> n </ci>
          </bvar>
          <apply>
            <times/>
            <ci> k </ci>
            <apply>
              <power/>
              <ci> A </ci>
              <apply>
                <plus/>
                <cn> 1 </cn>
```

```
                <ci> m </ci>
              </apply>
            </apply>
            <apply>
              <plus/>
              <cn> 1 </cn>
              <apply>
                <power/>
                <apply>
                  <divide/>
                  <ci> w </ci>
                  <ci> wk </ci>
                </apply>
                <ci> n </ci>
              </apply>
            </apply>
          </apply>
        </lambda>
      </math>
    </functionDefinition>
    <functionDefinition metaid="COPASI38" id="f_prod_lc3"
name="f_prod_lc3">
      <annotation>
        <COPASI xmlns="http://www.copasi.org/static/sbml">
          <rdf:RDF xmlns:dcterms="http://purl.org/dc/terms/"
xmlns:rdf="http://www.w3.org/1999/02/22-rdf-syntax-ns#">
            <rdf:Description rdf:about="#COPASI38">
              <dcterms:created>
                <rdf:Description>

<dcterms:W3CDTF>2017-11-23T14:50:31Z</dcterms:W3CDTF>
                </rdf:Description>
              </dcterms:created>
            </rdf:Description>
          </rdf:RDF>
        </COPASI>
      </annotation>
      <math xmlns="http://www.w3.org/1998/Math/MathML">
        <lambda>
          <bvar>
            <ci> k </ci>
          </bvar>
          <bvar>
            <ci> A </ci>
          </bvar>
          <bvar>
            <ci> K </ci>
          </bvar>
          <apply>
            <times/>
            <ci> k </ci>
            <apply>
```

```
                <divide/>
                <ci> A </ci>
                <apply>
                  <plus/>
                  <ci> K </ci>
                  <ci> A </ci>
                </apply>
              </apply>
            </apply>
          </lambda>
        </math>
      </functionDefinition>
    </listOfFunctionDefinitions>
    <listOfUnitDefinitions>
      <unitDefinition id="length" name="length">
        <listOfUnits>
          <unit kind="metre" exponent="1" scale="0" multiplier="1"/>
        </listOfUnits>
      </unitDefinition>
      <unitDefinition id="area" name="area">
        <listOfUnits>
          <unit kind="metre" exponent="2" scale="0" multiplier="1"/>
        </listOfUnits>
      </unitDefinition>
      <unitDefinition id="volume" name="volume">
        <listOfUnits>
          <unit kind="litre" exponent="1" scale="0" multiplier="1"/>
        </listOfUnits>
      </unitDefinition>
      <unitDefinition id="time" name="time">
        <listOfUnits>
          <unit kind="second" exponent="1" scale="0" multiplier="1"/>
        </listOfUnits>
      </unitDefinition>
      <unitDefinition id="substance" name="substance">
        <listOfUnits>
          <unit kind="item" exponent="1" scale="0" multiplier="1"/>
        </listOfUnits>
      </unitDefinition>
    </listOfUnitDefinitions>
    <listOfCompartments>
      <compartment metaid="COPASI1" id="ER" name="ER"
spatialDimensions="3" size="1" units="volume" constant="true">
        <annotation>
          <COPASI xmlns="http://www.copasi.org/static/sbml">
            <rdf:RDF xmlns:dcterms="http://purl.org/dc/terms/"
xmlns:rdf="http://www.w3.org/1999/02/22-rdf-syntax-ns#">
              <rdf:Description rdf:about="#COPASI1">
                <dcterms:created>
                  <rdf:Description>

<dcterms:W3CDTF>2017-11-22T13:32:41Z</dcterms:W3CDTF>
```

```
                  </rdf:Description>
                </dcterms:created>
              </rdf:Description>
            </rdf:RDF>
          </COPASI>
        </annotation>
      </compartment>
    </listOfCompartments>
    <listOfSpecies>
      <species metaid="COPASI2" id="ATG13" name="ATG13" compartment="ER"
initialConcentration="1" substanceUnits="substance"
hasOnlySubstanceUnits="false" boundaryCondition="false"
constant="false">
        <annotation>
          <COPASI xmlns="http://www.copasi.org/static/sbml">
            <rdf:RDF xmlns:dcterms="http://purl.org/dc/terms/"
xmlns:rdf="http://www.w3.org/1999/02/22-rdf-syntax-ns#">
              <rdf:Description rdf:about="#COPASI2">
                <dcterms:created>
                  <rdf:Description>

<dcterms:W3CDTF>2017-11-22T13:32:48Z</dcterms:W3CDTF>
                  </rdf:Description>
                </dcterms:created>
              </rdf:Description>
            </rdf:RDF>
          </COPASI>
        </annotation>
      </species>
      <species metaid="COPASI3" id="LC3" name="LC3" compartment="ER"
initialConcentration="0" substanceUnits="substance"
hasOnlySubstanceUnits="false" boundaryCondition="false"
constant="false">
        <annotation>
          <COPASI xmlns="http://www.copasi.org/static/sbml">
            <rdf:RDF xmlns:dcterms="http://purl.org/dc/terms/"
xmlns:rdf="http://www.w3.org/1999/02/22-rdf-syntax-ns#">
              <rdf:Description rdf:about="#COPASI3">
                <dcterms:created>
                  <rdf:Description>

<dcterms:W3CDTF>2017-11-23T14:50:55Z</dcterms:W3CDTF>
                  </rdf:Description>
                </dcterms:created>
              </rdf:Description>
            </rdf:RDF>
          </COPASI>
        </annotation>
      </species>
    </listOfSpecies>
    <listOfParameters>
      <parameter metaid="COPASI4" id="kprodATG13" name="kprodATG13"
```

```
value="0.0113491" constant="false">
        <annotation>
          <COPASI xmlns="http://www.copasi.org/static/sbml">
            <rdf:RDF xmlns:dcterms="http://purl.org/dc/terms/"
xmlns:rdf="http://www.w3.org/1999/02/22-rdf-syntax-ns#">
              <rdf:Description rdf:about="#COPASI4">
                <dcterms:created>
                  <rdf:Description>

<dcterms:W3CDTF>2016-09-20T16:37:39Z</dcterms:W3CDTF>
                  </rdf:Description>
                </dcterms:created>
              </rdf:Description>
            </rdf:RDF>
          </COPASI>
        </annotation>
      </parameter>
      <parameter metaid="COPASI5" id="ATG13_obs" name="ATG13_obs"
value="1111.7077740128" constant="false">
        <annotation>
          <COPASI xmlns="http://www.copasi.org/static/sbml">
            <rdf:RDF xmlns:dcterms="http://purl.org/dc/terms/"
xmlns:rdf="http://www.w3.org/1999/02/22-rdf-syntax-ns#">
              <rdf:Description rdf:about="#COPASI5">
                <dcterms:created>
                  <rdf:Description>

<dcterms:W3CDTF>2016-09-22T10:52:54Z</dcterms:W3CDTF>
                  </rdf:Description>
                </dcterms:created>
              </rdf:Description>
            </rdf:RDF>
          </COPASI>
        </annotation>
      </parameter>
      <parameter metaid="COPASI6" id="wrtm" name="wrtm" value="0"
constant="false">
        <annotation>
          <COPASI xmlns="http://www.copasi.org/static/sbml">
            <rdf:RDF xmlns:dcterms="http://purl.org/dc/terms/"
xmlns:rdf="http://www.w3.org/1999/02/22-rdf-syntax-ns#">
              <rdf:Description rdf:about="#COPASI6">
                <dcterms:created>
                  <rdf:Description>

<dcterms:W3CDTF>2016-09-22T15:46:47Z</dcterms:W3CDTF>
                  </rdf:Description>
                </dcterms:created>
              </rdf:Description>
            </rdf:RDF>
          </COPASI>
        </annotation>
```

```
      </parameter>
      <parameter metaid="COPASI7" id="wrtm_flag" name="wrtm_flag"
value="0" constant="true">
        <annotation>
          <COPASI xmlns="http://www.copasi.org/static/sbml">
            <rdf:RDF xmlns:dcterms="http://purl.org/dc/terms/"
xmlns:rdf="http://www.w3.org/1999/02/22-rdf-syntax-ns#">
              <rdf:Description rdf:about="#COPASI7">
                <dcterms:created>
                  <rdf:Description>

<dcterms:W3CDTF>2016-09-22T15:46:39Z</dcterms:W3CDTF>
                  </rdf:Description>
                </dcterms:created>
              </rdf:Description>
            </rdf:RDF>
          </COPASI>
        </annotation>
      </parameter>
      <parameter metaid="COPASI8" id="kwrtm" name="kwrtm"
value="1.73235" constant="true">
        <annotation>
          <COPASI xmlns="http://www.copasi.org/static/sbml">
            <rdf:RDF xmlns:dcterms="http://purl.org/dc/terms/"
xmlns:rdf="http://www.w3.org/1999/02/22-rdf-syntax-ns#">
              <rdf:Description rdf:about="#COPASI8">
                <dcterms:created>
                  <rdf:Description>

<dcterms:W3CDTF>2016-09-22T15:48:18Z</dcterms:W3CDTF>
                  </rdf:Description>
                </dcterms:created>
              </rdf:Description>
            </rdf:RDF>
          </COPASI>
        </annotation>
      </parameter>
      <parameter metaid="COPASI9" id="ATG13_sf" name="ATG13_sf"
value="1111.7077740128" constant="false">
        <annotation>
          <COPASI xmlns="http://www.copasi.org/static/sbml">
            <rdf:RDF xmlns:dcterms="http://purl.org/dc/terms/"
xmlns:rdf="http://www.w3.org/1999/02/22-rdf-syntax-ns#">
              <rdf:Description rdf:about="#COPASI9">
                <dcterms:created>
                  <rdf:Description>

<dcterms:W3CDTF>2016-09-27T15:55:01Z</dcterms:W3CDTF>
                  </rdf:Description>
                </dcterms:created>
              </rdf:Description>
            </rdf:RDF>
```

```
          </COPASI>
        </annotation>
      </parameter>
      <parameter metaid="COPASI10" id="ATG13_min" name="ATG13_min"
value="1111.7077740128" constant="true">
        <annotation>
          <COPASI xmlns="http://www.copasi.org/static/sbml">
            <rdf:RDF xmlns:dcterms="http://purl.org/dc/terms/"
xmlns:rdf="http://www.w3.org/1999/02/22-rdf-syntax-ns#">
              <rdf:Description rdf:about="#COPASI10">
                <dcterms:created>
                  <rdf:Description>

<dcterms:W3CDTF>2016-09-27T16:49:27Z</dcterms:W3CDTF>
                  </rdf:Description>
                </dcterms:created>
              </rdf:Description>
            </rdf:RDF>
          </COPASI>
        </annotation>
      </parameter>
      <parameter metaid="COPASI11" id="kremATG13" name="kremATG13"
value="0.0113493" constant="false">
        <annotation>
          <COPASI xmlns="http://www.copasi.org/static/sbml">
            <rdf:RDF xmlns:dcterms="http://purl.org/dc/terms/"
xmlns:rdf="http://www.w3.org/1999/02/22-rdf-syntax-ns#">
              <rdf:Description rdf:about="#COPASI11">
                <dcterms:created>
                  <rdf:Description>

<dcterms:W3CDTF>2016-10-17T11:48:37Z</dcterms:W3CDTF>
                  </rdf:Description>
                </dcterms:created>
              </rdf:Description>
            </rdf:RDF>
          </COPASI>
        </annotation>
      </parameter>
      <parameter metaid="COPASI12" id="n" name="n" value="1"
constant="true">
        <annotation>
          <COPASI xmlns="http://www.copasi.org/static/sbml">
            <rdf:RDF xmlns:dcterms="http://purl.org/dc/terms/"
xmlns:rdf="http://www.w3.org/1999/02/22-rdf-syntax-ns#">
              <rdf:Description rdf:about="#COPASI12">
                <dcterms:created>
                  <rdf:Description>

<dcterms:W3CDTF>2016-10-21T14:26:01Z</dcterms:W3CDTF>
                  </rdf:Description>
                </dcterms:created>
```

```
              </rdf:Description>
            </rdf:RDF>
          </COPASI>
        </annotation>
      </parameter>
      <parameter metaid="COPASI13" id="t" name="t"
value="38.4190105720273" constant="false">
        <annotation>
          <COPASI xmlns="http://www.copasi.org/static/sbml">
            <rdf:RDF xmlns:dcterms="http://purl.org/dc/terms/"
xmlns:rdf="http://www.w3.org/1999/02/22-rdf-syntax-ns#">
              <rdf:Description rdf:about="#COPASI13">
                <dcterms:created>
                  <rdf:Description>

<dcterms:W3CDTF>2016-10-21T16:51:48Z</dcterms:W3CDTF>
                  </rdf:Description>
                </dcterms:created>
              </rdf:Description>
            </rdf:RDF>
          </COPASI>
        </annotation>
      </parameter>
      <parameter metaid="COPASI14" id="wrtm_si" name="wrtm_si"
value="1500" constant="true">
        <annotation>
          <COPASI xmlns="http://www.copasi.org/static/sbml">
            <rdf:RDF xmlns:dcterms="http://purl.org/dc/terms/"
xmlns:rdf="http://www.w3.org/1999/02/22-rdf-syntax-ns#">
              <rdf:Description rdf:about="#COPASI14">
                <dcterms:created>
                  <rdf:Description>

<dcterms:W3CDTF>2016-10-26T16:07:58Z</dcterms:W3CDTF>
                  </rdf:Description>
                </dcterms:created>
              </rdf:Description>
            </rdf:RDF>
          </COPASI>
        </annotation>
      </parameter>
      <parameter metaid="COPASI15" id="wrtm_sf" name="wrtm_sf"
value="1500" constant="true">
        <annotation>
          <COPASI xmlns="http://www.copasi.org/static/sbml">
            <rdf:RDF xmlns:dcterms="http://purl.org/dc/terms/"
xmlns:rdf="http://www.w3.org/1999/02/22-rdf-syntax-ns#">
              <rdf:Description rdf:about="#COPASI15">
                <dcterms:created>
                  <rdf:Description>

<dcterms:W3CDTF>2016-10-26T16:08:06Z</dcterms:W3CDTF>
```

```
                </rdf:Description>
              </dcterms:created>
            </rdf:Description>
          </rdf:RDF>
        </COPASI>
      </annotation>
    </parameter>
    <parameter metaid="COPASI16" id="wrtm_scaled" name="wrtm_scaled"
value="0" constant="false">
      <annotation>
        <COPASI xmlns="http://www.copasi.org/static/sbml">
          <rdf:RDF xmlns:dcterms="http://purl.org/dc/terms/"
xmlns:rdf="http://www.w3.org/1999/02/22-rdf-syntax-ns#">
            <rdf:Description rdf:about="#COPASI16">
              <dcterms:created>
                <rdf:Description>

<dcterms:W3CDTF>2016-10-26T16:08:18Z</dcterms:W3CDTF>
                </rdf:Description>
              </dcterms:created>
            </rdf:Description>
          </rdf:RDF>
        </COPASI>
      </annotation>
    </parameter>
    <parameter metaid="COPASI17" id="m" name="m" value="1.01365"
constant="true">
      <annotation>
        <COPASI xmlns="http://www.copasi.org/static/sbml">
          <rdf:RDF xmlns:dcterms="http://purl.org/dc/terms/"
xmlns:rdf="http://www.w3.org/1999/02/22-rdf-syntax-ns#">
            <rdf:Description rdf:about="#COPASI17">
              <dcterms:created>
                <rdf:Description>

<dcterms:W3CDTF>2016-10-27T12:48:31Z</dcterms:W3CDTF>
                </rdf:Description>
              </dcterms:created>
            </rdf:Description>
          </rdf:RDF>
        </COPASI>
      </annotation>
    </parameter>
    <parameter metaid="COPASI18" id="ATG13_max" name="ATG13_max"
value="1779.3120503268" constant="true">
      <annotation>
        <COPASI xmlns="http://www.copasi.org/static/sbml">
          <rdf:RDF xmlns:dcterms="http://purl.org/dc/terms/"
xmlns:rdf="http://www.w3.org/1999/02/22-rdf-syntax-ns#">
            <rdf:Description rdf:about="#COPASI18">
              <dcterms:created>
                <rdf:Description>
```

```
<dcterms:W3CDTF>2017-11-23T14:43:04Z</dcterms:W3CDTF>
                  </rdf:Description>
                </dcterms:created>
              </rdf:Description>
            </rdf:RDF>
          </COPASI>
        </annotation>
      </parameter>
      <parameter metaid="COPASI19" id="EC50ATG13" name="EC50ATG13"
value="333.802138157" constant="false">
        <annotation>
          <COPASI xmlns="http://www.copasi.org/static/sbml">
            <rdf:RDF xmlns:dcterms="http://purl.org/dc/terms/"
xmlns:rdf="http://www.w3.org/1999/02/22-rdf-syntax-ns#">
              <rdf:Description rdf:about="#COPASI19">
                <dcterms:created>
                  <rdf:Description>

<dcterms:W3CDTF>2017-11-23T14:43:46Z</dcterms:W3CDTF>
                  </rdf:Description>
                </dcterms:created>
              </rdf:Description>
            </rdf:RDF>
          </COPASI>
        </annotation>
      </parameter>
      <parameter metaid="COPASI20" id="kpeak" name="kpeak"
value="136.137" constant="true">
        <annotation>
          <COPASI xmlns="http://www.copasi.org/static/sbml">
            <rdf:RDF xmlns:dcterms="http://purl.org/dc/terms/"
xmlns:rdf="http://www.w3.org/1999/02/22-rdf-syntax-ns#">
              <rdf:Description rdf:about="#COPASI20">
                <dcterms:created>
                  <rdf:Description>

<dcterms:W3CDTF>2017-11-23T14:46:32Z</dcterms:W3CDTF>
                  </rdf:Description>
                </dcterms:created>
              </rdf:Description>
            </rdf:RDF>
          </COPASI>
        </annotation>
      </parameter>
      <parameter metaid="COPASI21" id="cf" name="cf" value="0"
constant="false">
        <annotation>
          <COPASI xmlns="http://www.copasi.org/static/sbml">
            <rdf:RDF xmlns:dcterms="http://purl.org/dc/terms/"
xmlns:rdf="http://www.w3.org/1999/02/22-rdf-syntax-ns#">
              <rdf:Description rdf:about="#COPASI21">
```

```
                <dcterms:created>
                  <rdf:Description>

<dcterms:W3CDTF>2017-11-23T14:46:51Z</dcterms:W3CDTF>
                  </rdf:Description>
                </dcterms:created>
              </rdf:Description>
            </rdf:RDF>
          </COPASI>
        </annotation>
      </parameter>
      <parameter metaid="COPASI22" id="kprodLC3" name="kprodLC3"
value="0.779693" constant="false">
        <annotation>
          <COPASI xmlns="http://www.copasi.org/static/sbml">
            <rdf:RDF xmlns:dcterms="http://purl.org/dc/terms/"
xmlns:rdf="http://www.w3.org/1999/02/22-rdf-syntax-ns#">
              <rdf:Description rdf:about="#COPASI22">
                <dcterms:created>
                  <rdf:Description>

<dcterms:W3CDTF>2017-11-23T14:48:03Z</dcterms:W3CDTF>
                  </rdf:Description>
                </dcterms:created>
              </rdf:Description>
            </rdf:RDF>
          </COPASI>
        </annotation>
      </parameter>
      <parameter metaid="COPASI23" id="peak_delay_obs"
name="peak_delay_obs" value="0" constant="false">
        <annotation>
          <COPASI xmlns="http://www.copasi.org/static/sbml">
            <rdf:RDF xmlns:dcterms="http://purl.org/dc/terms/"
xmlns:rdf="http://www.w3.org/1999/02/22-rdf-syntax-ns#">
              <rdf:Description rdf:about="#COPASI23">
                <dcterms:created>
                  <rdf:Description>

<dcterms:W3CDTF>2017-11-23T14:54:35Z</dcterms:W3CDTF>
                  </rdf:Description>
                </dcterms:created>
              </rdf:Description>
            </rdf:RDF>
          </COPASI>
        </annotation>
      </parameter>
      <parameter metaid="COPASI24" id="time_fire" name="time_fire"
value="0" constant="false">
        <annotation>
          <COPASI xmlns="http://www.copasi.org/static/sbml">
            <rdf:RDF xmlns:dcterms="http://purl.org/dc/terms/"
```

```
xmlns:rdf="http://www.w3.org/1999/02/22-rdf-syntax-ns#">
              <rdf:Description rdf:about="#COPASI24">
                <dcterms:created>
                  <rdf:Description>

<dcterms:W3CDTF>2017-11-24T15:09:10Z</dcterms:W3CDTF>
                  </rdf:Description>
                </dcterms:created>
              </rdf:Description>
            </rdf:RDF>
          </COPASI>
        </annotation>
      </parameter>
      <parameter metaid="COPASI25" id="MT_surf" name="MT_surf"
value="2.82258980823599" constant="true">
        <annotation>
          <COPASI xmlns="http://www.copasi.org/static/sbml">
            <rdf:RDF xmlns:dcterms="http://purl.org/dc/terms/"
xmlns:rdf="http://www.w3.org/1999/02/22-rdf-syntax-ns#">
              <rdf:Description rdf:about="#COPASI25">
                <dcterms:created>
                  <rdf:Description>

<dcterms:W3CDTF>2017-12-06T10:25:38Z</dcterms:W3CDTF>
                  </rdf:Description>
                </dcterms:created>
              </rdf:Description>
            </rdf:RDF>
          </COPASI>
        </annotation>
      </parameter>
      <parameter metaid="COPASI26" id="MT_diam" name="MT_diam"
value="0.947870371202267" constant="true">
        <annotation>
          <COPASI xmlns="http://www.copasi.org/static/sbml">
            <rdf:RDF xmlns:dcterms="http://purl.org/dc/terms/"
xmlns:rdf="http://www.w3.org/1999/02/22-rdf-syntax-ns#">
              <rdf:Description rdf:about="#COPASI26">
                <dcterms:created>
                  <rdf:Description>

<dcterms:W3CDTF>2017-12-06T10:25:38Z</dcterms:W3CDTF>
                  </rdf:Description>
                </dcterms:created>
              </rdf:Description>
            </rdf:RDF>
          </COPASI>
        </annotation>
      </parameter>
      <parameter metaid="COPASI27" id="p" name="p" value="2.78977"
constant="true">
        <annotation>
```

```
          <COPASI xmlns="http://www.copasi.org/static/sbml">
            <rdf:RDF xmlns:dcterms="http://purl.org/dc/terms/"
xmlns:rdf="http://www.w3.org/1999/02/22-rdf-syntax-ns#">
              <rdf:Description rdf:about="#COPASI27">
                <dcterms:created>
                  <rdf:Description>

<dcterms:W3CDTF>2017-11-23T14:46:32Z</dcterms:W3CDTF>
                  </rdf:Description>
                </dcterms:created>
              </rdf:Description>
            </rdf:RDF>
          </COPASI>
        </annotation>
      </parameter>
      <parameter metaid="COPASI28" id="peak_num" name="peak_num"
value="0" constant="false">
        <annotation>
          <COPASI xmlns="http://www.copasi.org/static/sbml">
            <rdf:RDF xmlns:dcterms="http://purl.org/dc/terms/"
xmlns:rdf="http://www.w3.org/1999/02/22-rdf-syntax-ns#">
              <rdf:Description rdf:about="#COPASI28">
                <dcterms:created>
                  <rdf:Description>

<dcterms:W3CDTF>2018-01-09T09:11:50Z</dcterms:W3CDTF>
                  </rdf:Description>
                </dcterms:created>
              </rdf:Description>
            </rdf:RDF>
          </COPASI>
        </annotation>
      </parameter>
      <parameter id="ModelValue_0" name="Initial for kprodATG13"
value="0.0113491" constant="true">
        <annotation>
          <initialValue xmlns="http://copasi.org/initialValue"
parent="kprodATG13"/>
        </annotation>
      </parameter>
      <parameter id="ModelValue_18" name="Initial for kprodLC3"
value="0.779693" constant="true">
        <annotation>
          <initialValue xmlns="http://copasi.org/initialValue"
parent="kprodLC3"/>
        </annotation>
      </parameter>
      <parameter id="ModelValue_7" name="Initial for kremATG13"
value="0.0113493" constant="true">
        <annotation>
          <initialValue xmlns="http://copasi.org/initialValue"
parent="kremATG13"/>
```

```
      </annotation>
    </parameter>
  </listOfParameters>
  <listOfInitialAssignments>
    <initialAssignment symbol="ATG13">
      <math xmlns="http://www.w3.org/1998/Math/MathML">
        <apply>
          <divide/>
          <ci> ATG13_min </ci>
          <ci> ATG13_sf </ci>
        </apply>
      </math>
    </initialAssignment>
    <initialAssignment symbol="ATG13_min">
      <math xmlns="http://www.w3.org/1998/Math/MathML">
        <cn> 1111.7077740128 </cn>
      </math>
    </initialAssignment>
    <initialAssignment symbol="t">
      <math xmlns="http://www.w3.org/1998/Math/MathML">
        <apply>
          <ci> RNORMAL </ci>
          <cn> 38.33333333 </cn>
          <cn> 11.6904519445 </cn>
        </apply>
      </math>
    </initialAssignment>
    <initialAssignment symbol="ATG13_max">
      <math xmlns="http://www.w3.org/1998/Math/MathML">
        <cn> 1779.3120503268 </cn>
      </math>
    </initialAssignment>
    <initialAssignment symbol="MT_surf">
      <math xmlns="http://www.w3.org/1998/Math/MathML">
        <apply>
          <times/>
          <pi/>
          <apply>
            <power/>
            <ci> MT_diam </ci>
            <cn> 2 </cn>
          </apply>
        </apply>
      </math>
    </initialAssignment>
    <initialAssignment symbol="MT_diam">
      <math xmlns="http://www.w3.org/1998/Math/MathML">
        <apply>
          <ci> RNORMAL </ci>
          <cn> 0.7285371148 </cn>
          <cn> 0.103021783 </cn>
        </apply>
```

```
      </math>
    </initialAssignment>
    <initialAssignment symbol="ModelValue_0">
      <math xmlns="http://www.w3.org/1998/Math/MathML">
        <ci> kprodATG13 </ci>
      </math>
    </initialAssignment>
    <initialAssignment symbol="ModelValue_18">
      <math xmlns="http://www.w3.org/1998/Math/MathML">
        <ci> kprodLC3 </ci>
      </math>
    </initialAssignment>
    <initialAssignment symbol="ModelValue_7">
      <math xmlns="http://www.w3.org/1998/Math/MathML">
        <ci> kremATG13 </ci>
      </math>
    </initialAssignment>
  </listOfInitialAssignments>
  <listOfRules>
    <assignmentRule variable="wrtm">
      <math xmlns="http://www.w3.org/1998/Math/MathML">
        <piecewise>
          <piece>
            <apply>
              <divide/>
              <cn> 0 </cn>
              <ci> wrtm_sf </ci>
            </apply>
            <apply>
              <eq/>
              <ci> wrtm_flag </ci>
              <cn> 0 </cn>
            </apply>
          </piece>
          <otherwise>
            <apply>
              <divide/>
              <ci> wrtm_si </ci>
              <ci> wrtm_sf </ci>
            </apply>
          </otherwise>
        </piecewise>
      </math>
    </assignmentRule>
    <assignmentRule variable="ATG13_sf">
      <math xmlns="http://www.w3.org/1998/Math/MathML">
        <ci> ATG13_min </ci>
      </math>
    </assignmentRule>
    <assignmentRule variable="ATG13_obs">
      <math xmlns="http://www.w3.org/1998/Math/MathML">
        <apply>
```

```
        <times/>
        <ci> ATG13_sf </ci>
        <ci> ATG13 </ci>
      </apply>
    </math>
  </assignmentRule>
  <assignmentRule variable="wrtm_scaled">
    <math xmlns="http://www.w3.org/1998/Math/MathML">
      <apply>
        <times/>
        <ci> wrtm_sf </ci>
        <ci> wrtm </ci>
      </apply>
    </math>
  </assignmentRule>
  <assignmentRule variable="EC50ATG13">
    <math xmlns="http://www.w3.org/1998/Math/MathML">
      <apply>
        <divide/>
        <apply>
          <minus/>
          <ci> ATG13_max </ci>
          <ci> ATG13_min </ci>
        </apply>
        <cn> 2 </cn>
      </apply>
    </math>
  </assignmentRule>
  <assignmentRule variable="cf">
    <math xmlns="http://www.w3.org/1998/Math/MathML">
      <apply>
        <divide/>
        <ci> LC3 </ci>
        <ci> MT_surf </ci>
      </apply>
    </math>
  </assignmentRule>
  <assignmentRule variable="peak_delay_obs">
    <math xmlns="http://www.w3.org/1998/Math/MathML">
      <apply>
        <times/>
        <ci> kpeak </ci>
        <apply>
          <power/>
          <ci> cf </ci>
          <ci> p </ci>
        </apply>
      </apply>
    </math>
  </assignmentRule>
</listOfRules>
<listOfReactions>
```

```
      <reaction metaid="COPASI33" id="ATG13_prod" name="ATG13_prod"
reversible="false" fast="false">
        <annotation>
          <COPASI xmlns="http://www.copasi.org/static/sbml">
            <rdf:RDF xmlns:dcterms="http://purl.org/dc/terms/"
xmlns:rdf="http://www.w3.org/1999/02/22-rdf-syntax-ns#">
              <rdf:Description rdf:about="#COPASI33">
                <dcterms:created>
                  <rdf:Description>

<dcterms:W3CDTF>2017-11-22T13:38:36Z</dcterms:W3CDTF>
                  </rdf:Description>
                </dcterms:created>
              </rdf:Description>
            </rdf:RDF>
          </COPASI>
        </annotation>
        <listOfProducts>
          <speciesReference species="ATG13" stoichiometry="1"
constant="true"/>
        </listOfProducts>
        <kineticLaw>
          <math xmlns="http://www.w3.org/1998/Math/MathML">
            <apply>
              <times/>
              <ci> ER </ci>
              <apply>
                <ci> f_prod_atg13 </ci>
                <ci> kprodATG13 </ci>
                <ci> ATG13 </ci>
                <ci> m </ci>
                <ci> wrtm </ci>
                <ci> kwrtm </ci>
                <ci> n </ci>
              </apply>
            </apply>
          </math>
        </kineticLaw>
      </reaction>
      <reaction metaid="COPASI34" id="ATG13_rem" name="ATG13_rem"
reversible="false" fast="false">
        <annotation>
          <COPASI xmlns="http://www.copasi.org/static/sbml">
            <rdf:RDF xmlns:dcterms="http://purl.org/dc/terms/"
xmlns:rdf="http://www.w3.org/1999/02/22-rdf-syntax-ns#">
              <rdf:Description rdf:about="#COPASI34">
                <dcterms:created>
                  <rdf:Description>

<dcterms:W3CDTF>2017-11-22T13:43:39Z</dcterms:W3CDTF>
                  </rdf:Description>
                </dcterms:created>
```

```
            </rdf:Description>
          </rdf:RDF>
        </COPASI>
      </annotation>
      <listOfReactants>
        <speciesReference species="ATG13" stoichiometry="1"
constant="true"/>
      </listOfReactants>
      <kineticLaw>
        <math xmlns="http://www.w3.org/1998/Math/MathML">
          <apply>
            <times/>
            <ci> ER </ci>
            <apply>
              <ci> f_rem_atg13 </ci>
              <ci> kremATG13 </ci>
              <ci> ATG13 </ci>
              <ci> m </ci>
              <ci> wrtm </ci>
              <ci> kwrtm </ci>
              <ci> n </ci>
            </apply>
          </apply>
        </math>
      </kineticLaw>
    </reaction>
    <reaction metaid="COPASI35" id="LC3_production"
name="LC3_production" reversible="false" fast="false">
      <annotation>
        <COPASI xmlns="http://www.copasi.org/static/sbml">
          <rdf:RDF xmlns:dcterms="http://purl.org/dc/terms/"
xmlns:rdf="http://www.w3.org/1999/02/22-rdf-syntax-ns#">
            <rdf:Description rdf:about="#COPASI35">
              <dcterms:created>
                <rdf:Description>

<dcterms:W3CDTF>2017-11-23T14:49:53Z</dcterms:W3CDTF>
                </rdf:Description>
              </dcterms:created>
            </rdf:Description>
          </rdf:RDF>
        </COPASI>
      </annotation>
      <listOfProducts>
        <speciesReference species="LC3" stoichiometry="1"
constant="true"/>
      </listOfProducts>
      <listOfModifiers>
        <modifierSpeciesReference species="ATG13"/>
      </listOfModifiers>
      <kineticLaw>
        <math xmlns="http://www.w3.org/1998/Math/MathML">
```

```
            <apply>
              <times/>
              <ci> ER </ci>
              <apply>
                <ci> f_prod_lc3 </ci>
                <ci> kprodLC3 </ci>
                <ci> ATG13 </ci>
                <ci> EC50ATG13 </ci>
              </apply>
            </apply>
          </math>
        </kineticLaw>
      </reaction>
    </listOfReactions>
    <listOfEvents>
      <event metaid="COPASI29" id="atg13_accumulation"
name="atg13_accumulation" useValuesFromTriggerTime="false">
        <annotation>
          <COPASI xmlns="http://www.copasi.org/static/sbml">
            <rdf:RDF xmlns:dcterms="http://purl.org/dc/terms/"
xmlns:rdf="http://www.w3.org/1999/02/22-rdf-syntax-ns#">
              <rdf:Description rdf:about="#COPASI29">
                <dcterms:created>
                  <rdf:Description>

<dcterms:W3CDTF>2017-11-22T13:22:45Z</dcterms:W3CDTF>
                  </rdf:Description>
                </dcterms:created>
              </rdf:Description>
            </rdf:RDF>
          </COPASI>
        </annotation>
        <trigger initialValue="false" persistent="true">
          <math xmlns="http://www.w3.org/1998/Math/MathML">
            <apply>
              <and/>
              <apply>
                <and/>
                <apply>
                  <leq/>
                  <ci> ATG13_obs </ci>
                  <ci> ATG13_min </ci>
                </apply>
                <apply>
                  <eq/>
                  <ci> kremATG13 </ci>
                  <cn> 0 </cn>
                </apply>
              </apply>
              <apply>
                <lt/>
                <ci> cf </ci>
```

```
              <cn> 1 </cn>
            </apply>
          </apply>
        </math>
      </trigger>
      <delay>
        <math xmlns="http://www.w3.org/1998/Math/MathML">
          <ci> peak_delay_obs </ci>
        </math>
      </delay>
      <listOfEventAssignments>
        <eventAssignment variable="kprodATG13">
          <math xmlns="http://www.w3.org/1998/Math/MathML">
            <ci> ModelValue_0 </ci>
          </math>
        </eventAssignment>
        <eventAssignment variable="kremATG13">
          <math xmlns="http://www.w3.org/1998/Math/MathML">
            <cn> 0 </cn>
          </math>
        </eventAssignment>
        <eventAssignment variable="t">
          <math xmlns="http://www.w3.org/1998/Math/MathML">
            <apply>
              <ci> RNORMAL </ci>
              <cn> 38.33333333 </cn>
              <apply>
                <minus/>
                <apply>
                  <plus/>
                  <cn> 11.6904519445 </cn>
                  <ci> kprodATG13 </ci>
                </apply>
                <ci> kprodATG13 </ci>
              </apply>
            </apply>
          </math>
        </eventAssignment>
        <eventAssignment variable="time_fire">
          <math xmlns="http://www.w3.org/1998/Math/MathML">
            <csymbol encoding="text"
definitionURL="http://www.sbml.org/sbml/symbols/time"> time </csymbol>
          </math>
        </eventAssignment>
        <eventAssignment variable="kprodLC3">
          <math xmlns="http://www.w3.org/1998/Math/MathML">
            <ci> ModelValue_18 </ci>
          </math>
        </eventAssignment>
      </listOfEventAssignments>
    </event>
    <event metaid="COPASI30" id="atg13_removal" name="atg13_removal"
```

```
useValuesFromTriggerTime="false">
        <annotation>
          <COPASI xmlns="http://www.copasi.org/static/sbml">
            <rdf:RDF xmlns:dcterms="http://purl.org/dc/terms/"
xmlns:rdf="http://www.w3.org/1999/02/22-rdf-syntax-ns#">
              <rdf:Description rdf:about="#COPASI30">
                <dcterms:created>
                  <rdf:Description>

<dcterms:W3CDTF>2017-11-22T13:22:45Z</dcterms:W3CDTF>
                  </rdf:Description>
                </dcterms:created>
              </rdf:Description>
            </rdf:RDF>
          </COPASI>
        </annotation>
        <trigger initialValue="false" persistent="true">
          <math xmlns="http://www.w3.org/1998/Math/MathML">
            <apply>
              <and/>
              <apply>
                <gt/>
                <ci> kprodATG13 </ci>
                <cn> 0 </cn>
              </apply>
              <apply>
                <geq/>
                <csymbol encoding="text"
definitionURL="http://www.sbml.org/sbml/symbols/time"> time </csymbol>
                <apply>
                  <plus/>
                  <ci> time_fire </ci>
                  <ci> t </ci>
                </apply>
              </apply>
            </apply>
          </math>
        </trigger>
        <listOfEventAssignments>
          <eventAssignment variable="kprodATG13">
            <math xmlns="http://www.w3.org/1998/Math/MathML">
              <cn> 0 </cn>
            </math>
          </eventAssignment>
          <eventAssignment variable="kremATG13">
            <math xmlns="http://www.w3.org/1998/Math/MathML">
              <ci> ModelValue_7 </ci>
            </math>
          </eventAssignment>
          <eventAssignment variable="peak_num">
            <math xmlns="http://www.w3.org/1998/Math/MathML">
              <apply>
```

```
                &lt;plus/&gt;
                &lt;ci&gt; peak_num &lt;/ci&gt;
                &lt;cn&gt; 1 &lt;/cn&gt;
              &lt;/apply&gt;
            &lt;/math&gt;
          &lt;/eventAssignment&gt;
        &lt;/listOfEventAssignments&gt;
      &lt;/event&gt;
      <event metaid="COPASI31" id="atg13_basal" name="atg13_basal"
useValuesFromTriggerTime="false">
        &lt;annotation&gt;
          &lt;COPASI xmlns="http://www.copasi.org/static/sbml"&gt;
            <rdf:RDF xmlns:dcterms="http://purl.org/dc/terms/"
xmlns:rdf="http://www.w3.org/1999/02/22-rdf-syntax-ns#">
              &lt;rdf:Description rdf:about="#COPASI31"&gt;
                &lt;dcterms:created&gt;
                  &lt;rdf:Description&gt;

&lt;dcterms:W3CDTF&gt;2017-11-22T13:22:45Z&lt;/dcterms:W3CDTF&gt;
                  &lt;/rdf:Description&gt;
                &lt;/dcterms:created&gt;
              &lt;/rdf:Description&gt;
            &lt;/rdf:RDF&gt;
          &lt;/COPASI&gt;
        &lt;/annotation&gt;
        &lt;trigger initialValue="false" persistent="true"&gt;
          &lt;math xmlns="http://www.w3.org/1998/Math/MathML"&gt;
            &lt;apply&gt;
              &lt;and/&gt;
              &lt;apply&gt;
                &lt;leq/&gt;
                &lt;ci&gt; ATG13_obs &lt;/ci&gt;
                &lt;ci&gt; ATG13_min &lt;/ci&gt;
              &lt;/apply&gt;
              &lt;apply&gt;
                &lt;gt/&gt;
                &lt;ci&gt; kremATG13 &lt;/ci&gt;
                &lt;cn&gt; 0 &lt;/cn&gt;
              &lt;/apply&gt;
            &lt;/apply&gt;
          &lt;/math&gt;
        &lt;/trigger&gt;
        &lt;listOfEventAssignments&gt;
          &lt;eventAssignment variable="kprodATG13"&gt;
            &lt;math xmlns="http://www.w3.org/1998/Math/MathML"&gt;
              &lt;cn&gt; 0 &lt;/cn&gt;
            &lt;/math&gt;
          &lt;/eventAssignment&gt;
          &lt;eventAssignment variable="kremATG13"&gt;
            &lt;math xmlns="http://www.w3.org/1998/Math/MathML"&gt;
              &lt;cn&gt; 0 &lt;/cn&gt;
            &lt;/math&gt;
```

```
          </eventAssignment>
          <eventAssignment variable="kprodLC3">
            <math xmlns="http://www.w3.org/1998/Math/MathML">
              <cn> 0 </cn>
            </math>
          </eventAssignment>
          <eventAssignment variable="ATG13">
            <math xmlns="http://www.w3.org/1998/Math/MathML">
              <apply>
                <divide/>
                <ci> ATG13_min </ci>
                <ci> ATG13_sf </ci>
              </apply>
            </math>
          </eventAssignment>
        </listOfEventAssignments>
      </event>
      <event metaid="COPASI32" id="mt_engulfed" name="mt_engulfed"
useValuesFromTriggerTime="false">
        <annotation>
          <COPASI xmlns="http://www.copasi.org/static/sbml">
            <rdf:RDF xmlns:dcterms="http://purl.org/dc/terms/"
xmlns:rdf="http://www.w3.org/1999/02/22-rdf-syntax-ns#">
              <rdf:Description rdf:about="#COPASI32">
                <dcterms:created>
                  <rdf:Description>

<dcterms:W3CDTF>2017-11-22T13:22:45Z</dcterms:W3CDTF>
                  </rdf:Description>
                </dcterms:created>
              </rdf:Description>
            </rdf:RDF>
          </COPASI>
        </annotation>
        <trigger initialValue="false" persistent="true">
          <math xmlns="http://www.w3.org/1998/Math/MathML">
            <apply>
              <geq/>
              <ci> cf </ci>
              <cn> 1 </cn>
            </apply>
          </math>
        </trigger>
        <listOfEventAssignments>
          <eventAssignment variable="kprodATG13">
            <math xmlns="http://www.w3.org/1998/Math/MathML">
              <cn> 0 </cn>
            </math>
          </eventAssignment>
          <eventAssignment variable="kremATG13">
            <math xmlns="http://www.w3.org/1998/Math/MathML">
              <ci> ModelValue_7 </ci>
```

```
            </math>
          </eventAssignment>
          <eventAssignment variable="kprodLC3">
            <math xmlns="http://www.w3.org/1998/Math/MathML">
              <cn> 0 </cn>
            </math>
          </eventAssignment>
        </listOfEventAssignments>
      </event>
    </listOfEvents>
  </model>
</sbml>
```
